# IBEX – A versatile multi-plex optical imaging approach for deep phenotyping and spatial analysis of cells in complex tissues

**DOI:** 10.1101/2020.11.20.390690

**Authors:** Andrea J. Radtke, Evelyn Kandov, Bradley Lowekamp, Emily Speranza, Colin J. Chu, Anita Gola, Nishant Thakur, Rochelle Shih, Li Yao, Ziv Rafael Yaniv, Rebecca T. Beuschel, Juraj Kabat, Joshua Croteau, Jeremy Davis, Jonathan M. Hernandez, Ronald N. Germain

**Author notes:** Co-first authors (Andrea J. Radtke, Evelyn Kandov). Co-corresponding authors: Andrea Radtke; Ronald Germain. **Author Contributions** A.J.R and R.N.G. wrote the manuscript. A.J.R, E.K., B.L, C.J.C., E.S., A.G., N.T., R.S., Z.R.Y, R.T.B., and J.K designed, performed, and analyzed experiments. L.Y. and A.G. assisted with display items. J.C., J.D., and J.M.H. provided technical insight, reagents, and tissues. All authors helped write the manuscript.

## Abstract

The diverse composition of mammalian tissues poses challenges for understanding the cell-cell interactions required for organ homeostasis and how spatial relationships are perturbed during disease. Existing methods such as single-cell genomics, lacking a spatial context, and traditional immunofluorescence, capturing only 2-6 molecular features, cannot resolve these issues. Imaging technologies have been developed to address these problems, but each possesses limitations that constrain widespread use. Here we report a new method that overcomes major impediments to highly multi-plex tissue imaging. <u>I</u>terative <u>B</u>leaching <u>E</u>xtends multi-ple<u>X</u>ity (IBEX) uses an iterative staining and chemical bleaching method to enable high resolution imaging of >65 parameters in the same tissue section without physical degradation. IBEX can be employed with various types of conventional microscopes and permits use of both commercially available and user-generated antibodies in an ‘open’ system to allow easy adjustment of staining panels based on ongoing marker discovery efforts. We show how IBEX can also be used with amplified staining methods for imaging strongly fixed tissues with limited epitope retention and with oligonucleotide-based staining, allowing potential cross-referencing between flow cytometry, Cellular Indexing of Transcriptomes and Epitopes by Sequencing (CITE-Seq), and IBEX analysis of the same tissue. To facilitate data processing, we provide an open source platform for automated registration of iterative images. IBEX thus represents a technology that can be rapidly integrated into most current laboratory workflows to achieve high content imaging to reveal the complex cellular landscape of diverse organs and tissues.

**Significance Statement:** Single cell flow cytometry and genomic methods are rapidly increasing our knowledge of the diversity of cell types in metazoan tissues. However, suitably robust methods for placing these cells in a spatial context that reveal how their localization and putative interactions contribute to tissue physiology and pathology are still lacking. Here we provide a readily accessible pipeline (IBEX) for highly multi-plex immunofluorescent imaging that enables a fine-grained analysis of cells in their tissue context. Additionally, we describe extensions of the IBEX workflow to handle hard to image tissue preparations and a method to facilitate direct integration of the imaging data with flow cytometry and sequencing technologies.

## Introduction

Mammalian tissues are composed of a wide variety of cell types, presenting a major challenge to understanding the cell-cell interactions required for homeostasis as well as the compositional changes associated with disease. To address this complexity, several multi-plexed imaging methods utilizing conventional microscopes and commercially available antibodies have been described to overcome the target detection limitations of conventional immunohistochemistry (IHC) or immunofluorescence (IF) imaging (1–8). The majority of these methods generate high dimensional datasets through an iterative, multi-step process (a cycle) that includes: 1) immunolabeling with antibodies, 2) image acquisition, and 3) fluorophore inactivation or antibody/chromogen removal. While these methods are capable of generating high dimensional datasets, they are greatly limited by the number of markers visualized per cycle, length of time required for each cycle, or involve special fluid-handling platforms not generally available to most laboratories (1). Commercial systems based on the co-detection by indexing (CODEX) method (9) have facilitated the acquisition of multi-plex imaging data by providing a fully automated instrument for cyclic imaging. Despite this advancement, the proprietary nature of this method imposes constraints on the reagents available for use as well as the number of markers to be imaged for each round. Furthermore, cyclic imaging methods that employ a small number of markers per cycle (<3) may result in tissue loss due to the stress of repeated fluid exchanges. To this end, novel imaging techniques such as multi-plexed ion beam imaging (MIBI) (10) and imaging mass cytometry (IMC) (11) enable the capture of multi-parameter data without cyclic imaging. However, both of these methods require specialized instrumentation and consumables, with the latter often again limited in breadth to choices made by the supplier, not the investigator. This constrains their capacity for broadly analyzing human or experimental animal tissues with respect to lists of validated antibodies, the ability to work across various established protocols for tissue processing, and the capacity for real time changes to the epitope target list based on data emerging from high content methods such as single cell RNA sequencing (scRNA-Seq).

To facilitate the increasing need for high content analysis of tissues for projects such as the Human Cell Atlas and others, the field needs a fully open and extensible method for multi-plex imaging. Our laboratory has extensively characterized murine and human immune responses using quantitative multi-parameter imaging of fixed frozen samples (12–18). Importantly, this method of tissue fixation preserves tissue architecture and cellular morphology, is archivable, compatible with large volume imaging (19), and in its optimal form, eliminates technical challenges posed by formalin-fixed paraffin embedded (FFPE) samples. Leveraging this experience and our original single cycle histo-cytometry method for multi-plex data acquisition (12), we have now developed <u>I</u>terative <u>B</u>leaching <u>E</u>xtends multi-ple<u>X</u>ity (IBEX). This imaging technique reduces the time per cycle, uses a high number of antibodies per cycle, employs widely available reagents and instruments, provides open source software for image alignment, and minimizes physical damage to the tissue during multiple imaging cycles. Beyond the basic IBEX workflow, we have developed extensions to achieve multi-parameter imaging of heavily fixed tissues with limited retention of target epitopes and have incorporated commercially available oligonucleotide-conjugated antibodies to enable direct cross-comparisons to flow cytometry and scRNA-Seq data obtained by the Cellular Indexing of Transcriptomes and Epitopes by Sequencing (CITE-Seq) method (20). In addition to describing the specifics of the IBEX method, we provide multiple examples of the use of IBEX to analyze both immune and parenchymal cells in a diverse array of mouse and human tissues to illustrate the general applicability of the method. The IBEX method described here can be rapidly integrated into current laboratory workflows to obtain high dimensional imaging datasets of a wide range of animal and human tissues.

## Results

### IBEX builds and improves upon existing iterative imaging techniques

Iterative imaging methods typically use either fluorophore bleaching or antibody/chromogen removal to achieve multi-parameter datasets (1–8). Due to the harsh and variable conditions required to remove chromogens and antibodies with diverse target affinities, we pursued a strategy based on fluorophore bleaching. To achieve an efficient means to increase the number of markers visualized on a single section, we sought a fluorophore inactivation method that could bleach a wide range of fluorophores in minutes without epitope loss or tissue destruction. While H_2_O_2_ in alkaline solution has been reported to inactivate Cy3- and Cy5-conjugated antibodies in human FFPE samples (3), we observed significant tissue loss using this formulation over multiple cycles with fixed frozen samples (Fig. S1A). Adams *et al*. demonstrated the initial feasibility of borohydride derivates to bleach fluorophores; however, their fluorophore quenching method required 2 hours per cycle and comprised only 3 distinct imaging channels (6), making direct application for highly multi-plex imaging impractical. To expand upon this method, we tested antibodies directly conjugated to fluorophores with excitation and emission spectra spanning from 405 nm to 750 nm. We consistently found that the following fluorophores were inactivated within 15 minutes of exposure to 1 mg/ml of Lithium Borohydride (LiBH_4_): Pacific Blue, Alexa Fluor (AF)488, FITC, AF532, Phycoerythrin (PE), AF555, eFluor(eF)570, AF647, eF660, and AF700. Brilliant Violet conjugates BV421 and BV510 bleached within 15 minutes of exposure to 1 mg/ml of LiBH_4_ in the presence of light. In contrast, AF594, eF615, and the nuclear markers JOJO-1 and Hoechst required more than 120 minutes for significant loss of fluorescence signal (Table S1), permitting these probes to be used as fiducials for alignment of images emerging from iterative cycles.

To prevent tissue destruction over multiple cycles, we evaluated several different tissue adhesives and found that chrome gelatin alum securely adhered tissues to glass coverslips and slides, permitting more than 15 cycles to be performed with no appreciable loss to the tissue (Fig. S1B-C). We next reduced the antibody labeling time from 6-12 hours to 30-45 minutes by designing programs for a non-heating microwave that facilitates rapid antibody penetration into the section. Finally, although IBEX was designed to simply bleach the fluorophores, it was important to assess whether LiBH_4_ treatment physically removes antibodies from the tissue as this would have direct bearing on both the design and order of imaging panels. To examine this issue, mouse lymph node (LN) sections were immunolabeled with various primary antibodies, imaged, treated with LiBH_4_, and then incubated with a secondary antibody that would react with the primary antibody if it was still present on the tissue. For most of the antibody isotypes tested, we found almost identical staining patterns with the primary and secondary antibodies, indicating that LiBH_4_ acts primarily by fluorophore bleaching without stripping the fluorescently conjugated antibodies themselves (Fig. S2).

The resulting method, IBEX, reduces the fluorophore inactivation and antibody labeling steps to less than 1 hour (Fig. 1A). We first tested IBEX in practical use by examining the image quality that could be obtained from a 3 cycle analysis of mouse LNs (Fig. 1B), a tissue with which we had extensive experience using multi-parameter, single cycle staining and image collection. This initial test used 6-8 markers per cycle and showed that all fluorophores, except for AF594 and JOJO-1, bleached rapidly in the presence of LiBH_4_ treatment +/− light with no appreciable signal present after 10 minutes (Fig. 1C-D). These findings show that the IBEX pipeline performs as designed and allows for the rapid capture of high quality, multi-plexed imaging data over multiple cycles without tissue loss.

**Figure 1.**
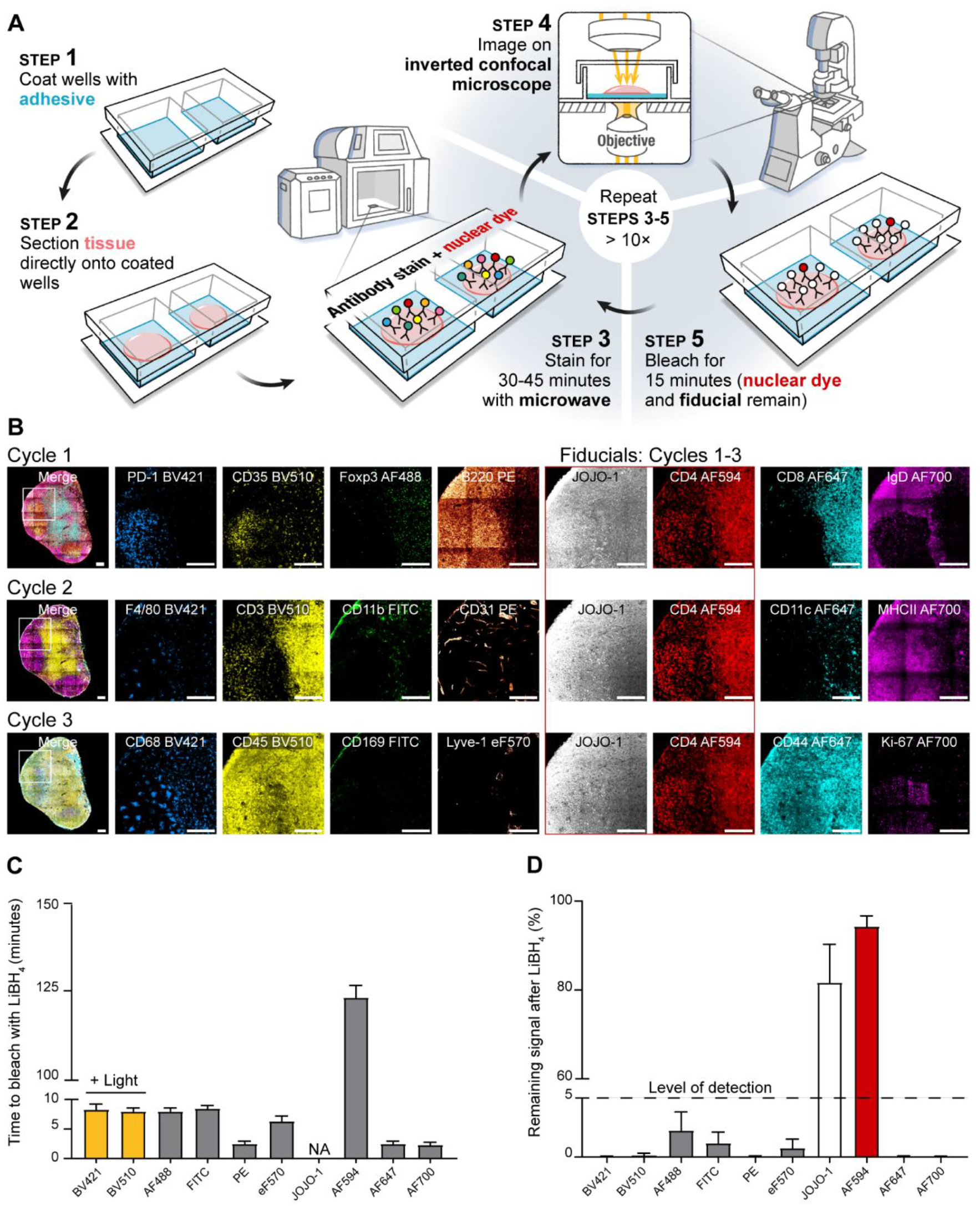
IBEX: A high dimensional, iterative imaging technique. *(A)* Schematic depicting IBEX protocol using an inverted confocal microscope. *(B)* Mice were immunized s.c. with 25 μl of SRBCs on day 0 and 7. On day 14, pLN tissue sections were labeled with 3 separate imaging panels. JOJO-1 and CD4 AF594 were present throughout cycles 1-3 and served as fiducials. Left-most panel is a composite of all channels except for JOJO-1. Scale bar represents 150 μm. Light refers to bleaching with LiBH_4_ while sample was illuminated with a metal halide lamp and DAPI filter *(C)* Time required to bleach respective fluorophores using LiBH_4_. NA indicates no appreciable loss of signal over multiple hours of LiBH_4_ exposure. *(D)* Percentage of fluorophore signal remaining after 15 minutes of LiBH_4_ treatment. Data are pooled from 2 similar experiments.

### A SimpleITK image registration pipeline

The IBEX method yields a series of images that are collected separately. To properly process these data and correctly assign markers to individual cells, it is essential that all the images be aligned at high resolution. While various registration algorithms have been reported (21, 22), we sought a method that could align large datasets, was flexible in terms of the repeated markers (fiducials) utilized, and provided both a qualitative and quantitative metric for registration. For these reasons, we developed a workflow using SimpleITK, a simplified open source interface to the Insight Toolkit (ITK) that is compatible with multiple programming languages (23, 24). The SimpleITK workflow is an image intensity-based form of image alignment that relies on a repeated marker channel (fiducial) for registration. Due to the resistance of AF594, JOJO-1, and Hoechst to bleaching, we utilized markers in these fluorophores as fiducials. For a multi-cycle IBEX experiment, a ‘fixed’ image z-stack was selected and all other ‘moving’ images were resampled to this image. A cross correlation matrix was generated on the repeated marker channels to provide a quantitative means for assessing the quality of image registration (Fig. 2A). To test the fidelity of this method, a 3 cycle IBEX experiment was performed on mouse spleen sections labeled with the nuclear marker JOJO-1 and membrane label CD4 AF594. For these experiments, JOJO-1 was selected as the fiducial used for image registration; however, CD4 (also repeated in these experiments) showed pixel-to-pixel alignment as reflected in the images (Fig. 2B). Importantly, this platform can readily scale to handle large datasets (>260 GB) comprised of 20 cycles of imaging (Fig. S1B-D). The SimpleITK registration workflow thus provides the needed cell-cell registration across x-y-z dimensions obtained via iterative imaging cycles using IBEX.

**Figure 2.**
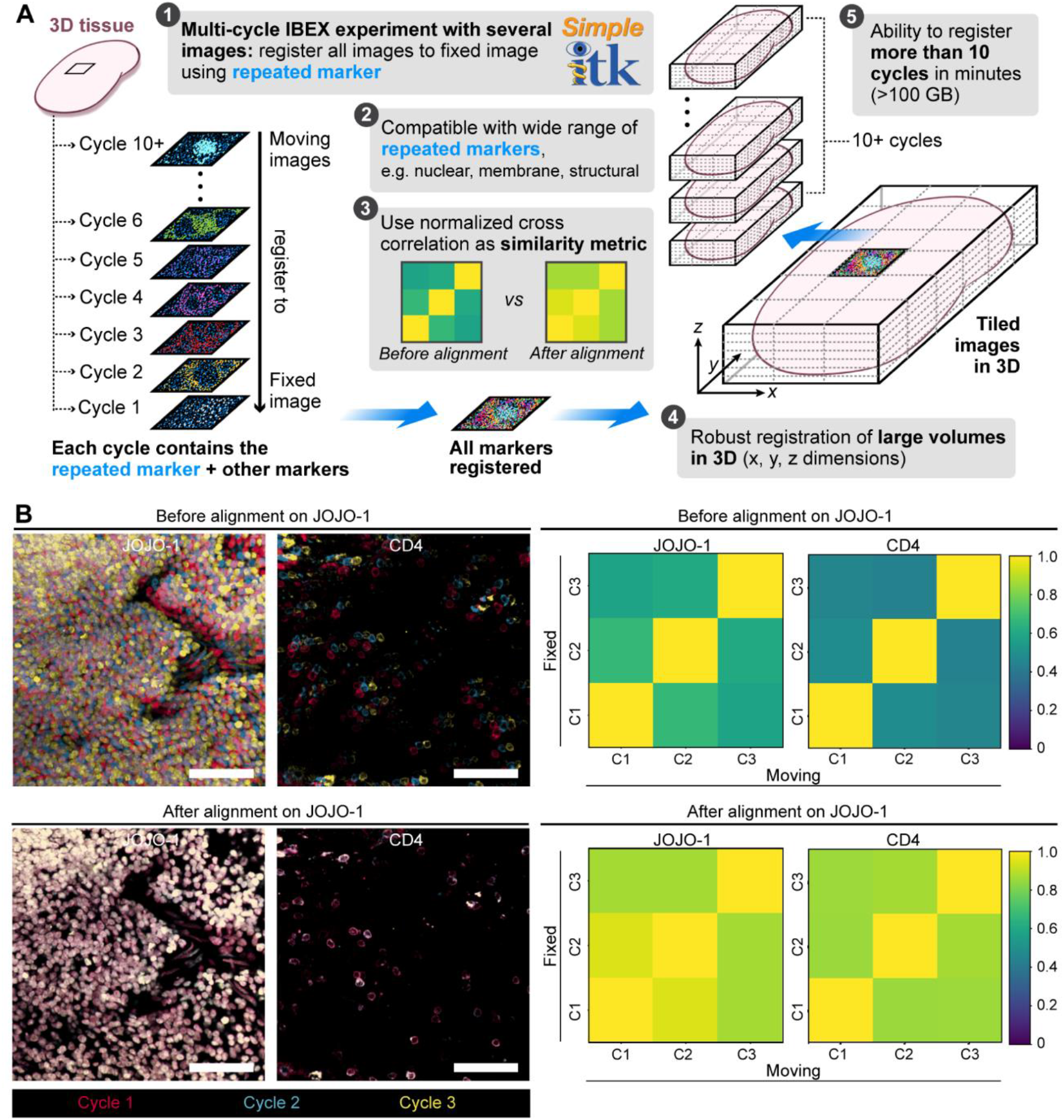
Image alignment with SimpleITK image registration pipeline. *(A)* Workflow for SimpleITK image registration pipeline. *(B)* Confocal images showing JOJO-1 and CD4 from 3 consecutive IBEX cycles before and after alignment using the nuclear marker JOJO-1 as a fidicual across all 3 cycles. CD4 was also repeated and shows cell-cell registration after JOJO-1 alignment. Cycle (C), scale bar is 50 μm. Cross correlation similarity matrices before and after alignment with JOJO-1 for JOJO-1 and CD4 channels. All experiments are representative of at least 2 similar experiments.

### IBEX is a versatile imaging method

One obstacle to the wide adoption of existing multi-plex imaging methods is the need for specialized instruments or custom imaging chambers, a luxury not afforded to all laboratories. In contrast, IBEX is easily adaptable to diverse microscope systems and has no restrictions on the substrate (slide, coverslip, etc.) used for imaging (Fig. S3A). As a proof-of-concept, immunized mouse LN sections adhered to slides were visualized using an upright confocal microscope (4 cycles, 6 markers per cycle) or an inverted fluorescence microscope (4 cycles, 4 markers per cycle) (Fig. S3B-C). These results demonstrate the compatibility of IBEX with a wide range of imaging systems; however, it is worth noting that the microscope system (confocal versus widefield) and configuration (light source, detectors, filter cubes) will dictate the image acquisition time, number of markers per cycle, and sample type that can be effectively imaged (5 vs. 30 μm tissue thickness).

In the case of animal studies, it is also very useful to be able to integrate antibody staining with imaging of fluorescent marker proteins expressed by engineered cells transferred into animals or expressed *in situ*. We therefore next investigated whether the IBEX method could be used to image tissues from animals expressing fluorescent proteins (25). To this end, high dimensional imaging was performed on thymic tissues from transgenic animals expressing the following fluorescent proteins (FPs): cyan (CFP), green (GFP), yellow (YFP), and red (RFP) (26). No appreciable loss in signal was observed after 10 IBEX cycles for any of the FPs examined (Fig. S4A-B). The photostable FPs were used as fiducials for a 4 cycle IBEX experiment that incorporated the bleachable fluorophores AF647 and AF700, yielding a dataset that provided information on clonality (CFP, GFP, YFP, RFP) of T cells (CD4, CD8, Foxp3) and myeloid cells (CD11c, MHC II) in the thymus (Fig. S4C).

To determine how IBEX performs using sections from a variety of tissues, we performed 3-5 cycle IBEX experiments on murine spleen, thymus, lung, small intestine, and liver tissue sections (Fig. 3A-B, Movies S1-S5, Table S2). It is important to note that the cycle and marker numbers described here are provided as a proof-of-concept and do not reflect technical limitations of the method. The antibody panels were designed to capture the major cellular populations and structures present in each organ and fluorophores were chosen to avoid native tissue autofluorescence. Organ-specific fiducials were selected based on expression throughout the tissue, e.g., EpCAM to mark the epithelium of the small intestine and laminin for the liver sinusoids. Collectively, these data confirm the ability to use IBEX to obtain high quality, multi-plexed imaging datasets from a wide range of tissues.

**Figure 3.**
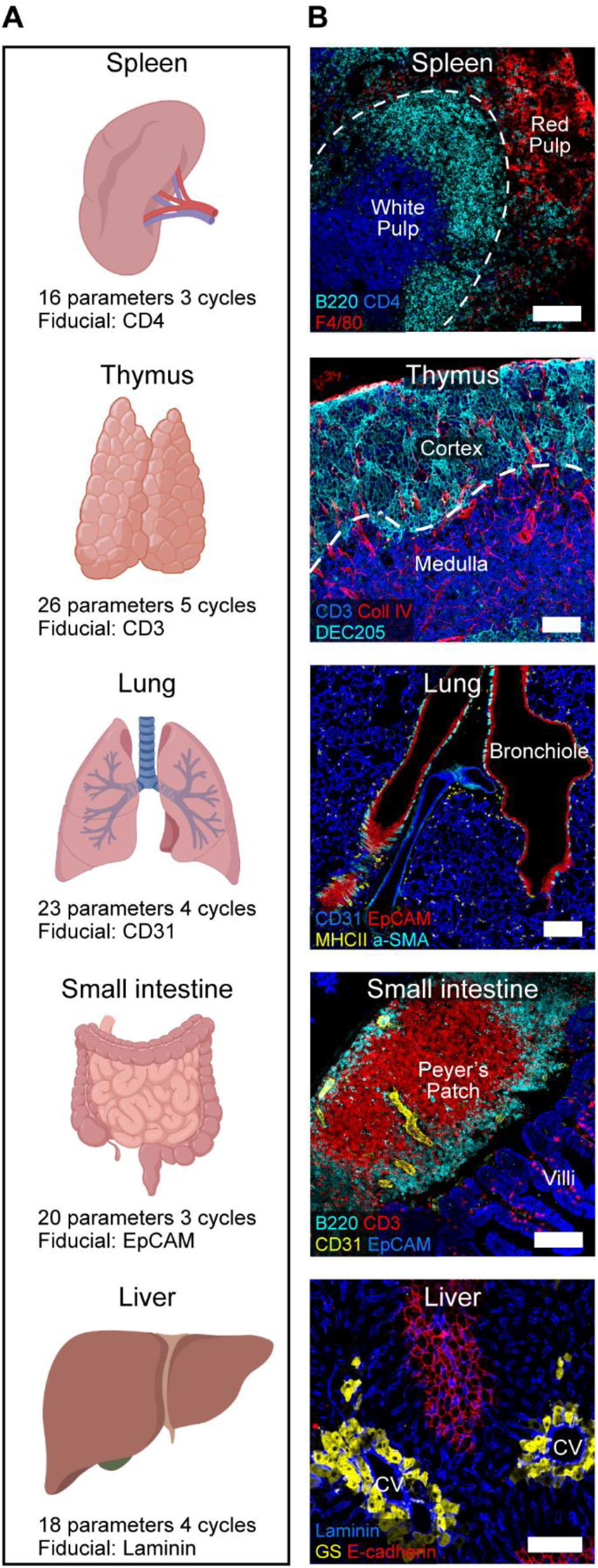
IBEX in multiple murine organs. *(A)* IBEX experimental parameters. (*B)* Confocal images from IBEX experiments in various mouse organs. Liver: central vein (CV) and glutamine synthetase (GS). Scale bar is 100 μm. See Movies S1-S5 for additional details.

### IBEX enables highly multi-plex, quantitative imaging

The design principles of IBEX were chosen to enable a very high number of parameters to be attained in the analysis of an individual tissue sample. To determine how extensively the multi-plexing capacity of IBEX can be pushed, we first performed 10 cycle 41 parameter IBEX experiments on LNs obtained from naïve and sheep red blood cell (SRBC)-immunized mice (Fig. 4A, Table S2, Movie S6). While epitope loss has been described for other iterative imaging techniques (5), we minimized this problem by increasing the number of markers per cycle and grouping markers present on the same cell in the same cycle. We observed qualitatively similar staining patterns when antibody panels were applied on individual sections alone (serial) versus on the same section iteratively (IBEX) (Fig. S5A and Movie S7). Therefore, quantitative differences observed between the two methods likely reflect biological differences resulting from variations in the magnitude of the immune response in individual LNs and not technical differences associated with epitope loss or steric hindrance (Fig. S5B and Movie S7).

**Figure 4.**
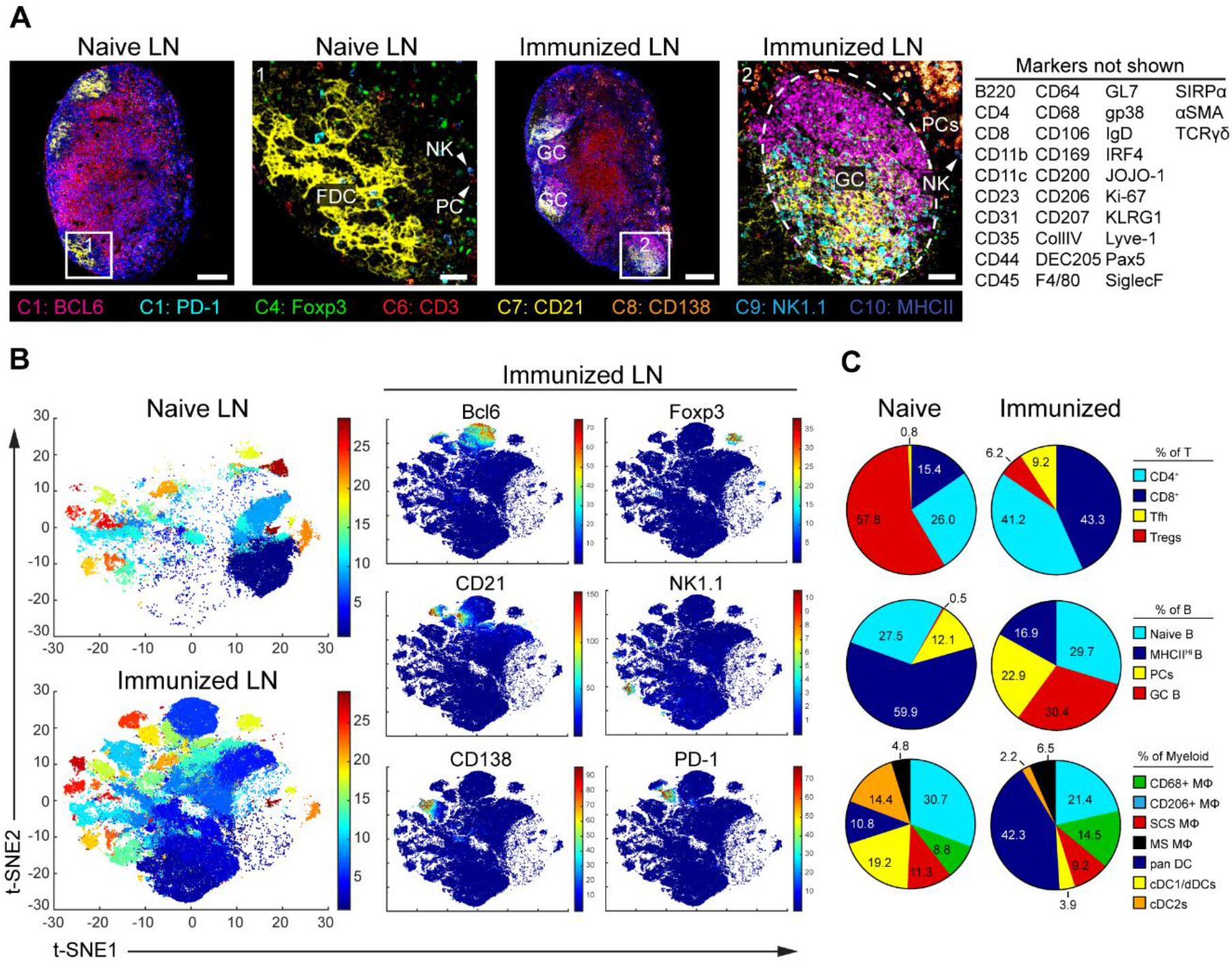
Visualization and quantification of LN populations using IBEX and histoCAT following immune perturbation. *(A)* Confocal images of pLNs from naïve and SRBC-immunized mice from 10 cycle (C) 41 parameter IBEX experiments, GC (germinal center), NK (natural killer), FDC (follicular dendritic cell), PC (plasma cells). Scale bars from left to right: 100 μm, 25 μm, 100 μm, and 50 μm. (*B)* t-SNE plots from naïve and immunized LNs identified by Phenograph clustering using segmented cells in histoCAT (naïve n = 32,091; immune n = 80,355). Color reflects the cluster ID number (1–29). Single plots show separation of representative markers into discrete clusters with color map showing relative expression levels based on Z-score normalized marker intensity values. *(C)* Phenograph clusters identified by histoCAT were phenotyped based on marker expression and expressed as a proportion of lineage. Tfh (T follicular helper), MΦ (macrophage), SCS (subcapsular sinus), MSM (Medullary sinus), DC (dendritic cell), dDC (dermal DC). Data are from one experiment and representative of 2 similar experiments. See Fig. S6 and Movie S6.

To assess the quality of data generated by the IBEX method, we employed the open source, computational histology topography cytometry analysis toolbox (histoCAT) to quantify differences in LN organization resulting from immunization (27). Individual cells were segmented based on membrane and nuclear labels with Ilastik (28) and CellProfiler (29) and then analyzed using the histoCAT graphical user interface (GUI) (Fig. S6A). The unsupervised clustering algorithm Phenograph (30) identified 29 phenotype clusters shared across the naïve and immunized LNs that could be visualized using the data dimensionality reduction method t-SNE (31) in histoCAT (Fig. 4B). As a testament to the fidelity of cell-cell alignment, phenotype clusters were often characterized by the expression of several different markers present in distinct imaging cycles (Fig. S6B-C). The abundance of these cell phenotypes varied from naïve and immunized LNs (Fig. S6D) and could be manually annotated based on marker expression to reveal an increase in plasma cells (PCs) (cluster 10) and germinal center (GC) B cells (cluster 6) in immunized LNs (Fig. 4C). As histoCAT relies on nuclear and membrane-based cell segmentation, it suffers from limitations frequently encountered with this approach: miscalling of phenotypes due to spatial overlap and improper segmentation of non-lymphocyte populations (32). The former is evident in the identification of the B cell specific transcription factor pax5 (33) on CD3^+^CD4^+^PD-1^+^Bcl6^+^ T follicular helper (Tfh) cells (cluster 19), an artifact due to the close proximity of these cells within the GC (Fig. S6C). Nevertheless, the results presented here demonstrate that IBEX-generated images are compatible with established methods for analyzing high dimensional imaging data.

### IBEX scales to capture ultra-high content data from large areas of human tissues

In addition to capturing the cellular landscape of a diverse range of murine tissues, the IBEX method scales to enable high content imaging of human tissues. To this end, we present an application of the IBEX method to visualize tumor-immune interactions in a human pancreatic LN with metastatic lesions as shown by CD138+EpCAM+ cells in the LN capsule and sinuses (Fig. 5A, Table S3). Interestingly, cancer cells were segregated from lymphocytes by extensive collagen remodeling and recruitment of myeloid cells expressing the secreted protein acidic and rich in cysteine (SPARC), a matricellular glycoprotein involved in extracellular matrix (ECM) deposition and implicated in metastasis (34). To move to even deeper analysis of a much larger sample, we used a total of 66 antibodies for the analysis of a > 3mm^2^ human mesenteric LN section. Using this approach, we were able to deeply phenotype a wide range of immune cells while observing no tissue degradation over 20 cycles (Fig. 5B, Movie S8, Table S3). Additionally, we observed subcellular resolution for PC-specific markers (membrane: CD138, nuclear: IRF4, cytoplasmic: IgA1, IgA2) present in distinct cycles with no epitope loss, as evidenced by our ability to label the immune marker CD45 with two different antibody clones present in cycles 9 and 19 (Movie S8). The utility of this method is further exemplified by our ability to characterize the complex stroma of the LN, shown to contain 9 distinct clusters in scRNA-Seq experiments from mouse LNs (35), using a wide range of antibodies visualized *in situ*: α-SMA, CD21, CD23, CD34, CD35, CD49a, CXCL12, CXCL13, desmin, fibronectin, and vimentin. These data highlight the capacity of IBEX to identify a large number of distinct cell types in clinically relevant samples, while also placing these components in a spatial setting missed by methods employing dissociated single cells.

**Figure 5.**
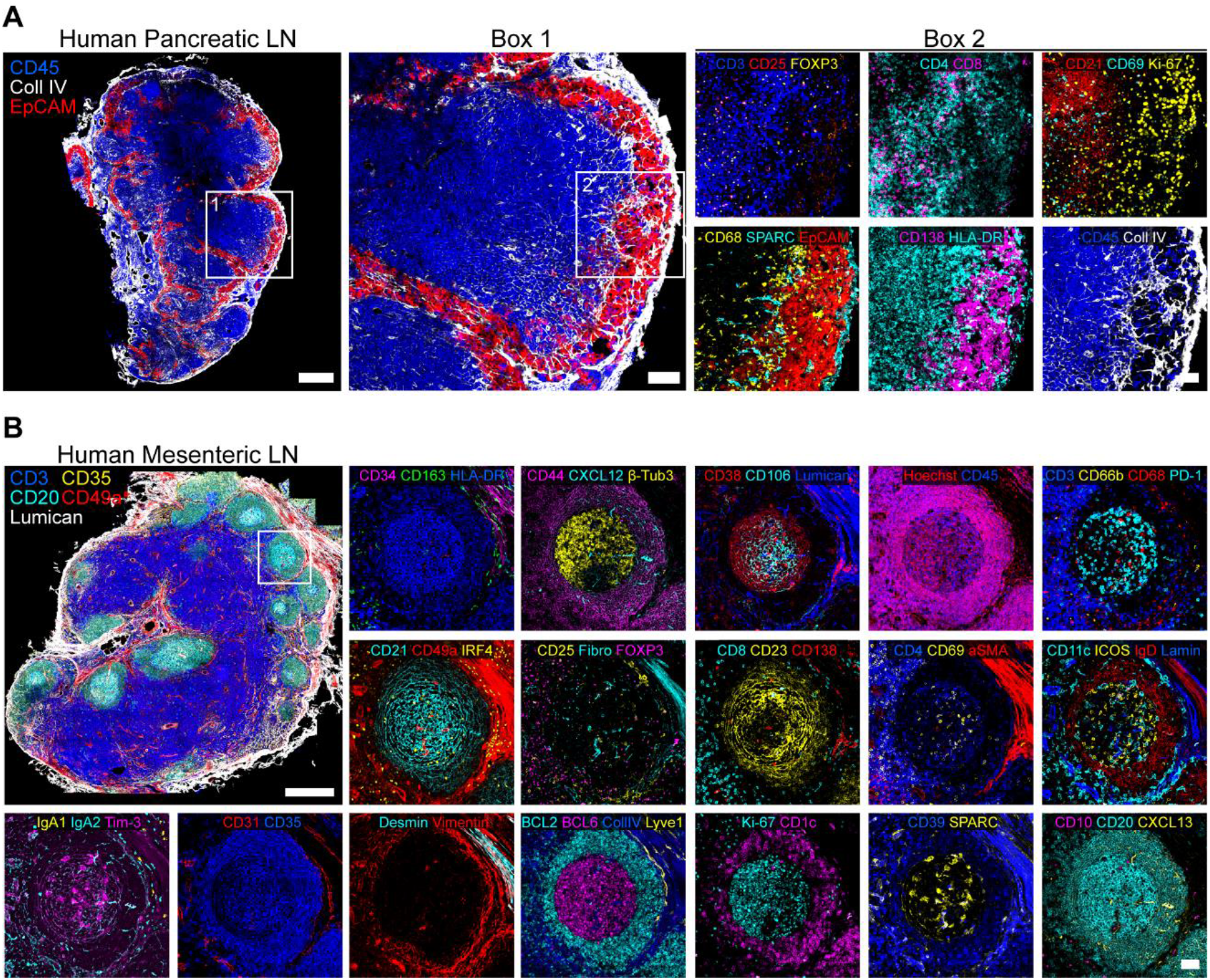
IBEX scales to capture ultra-high content imaging in human tissues. *(A)* Confocal images from a pancreatic LN with metastatic lesions from a patient with colorectal cancer (4 cycle 17 parameter IBEX experiment). Scale bar is 500 μm (left), 100 μm (Box 1), or 50 μm (Box 2). *(B)* Representative confocal images from human mesenteric LN obtained by IBEX method (20 cycle 66 parameters). Scale bars (500 or 50 μm). Fibronectin (Fibro), Laminin (Lamin). See Movie S8 for additional details.

### Extensions of IBEX workflow

Given the inherent versatility of the IBEX method, we sought to extend our workflow to develop two unique protocols, one that enables detection of low abundance epitopes and another that permits iterative imaging with oligonucleotide-conjugated antibodies corresponding to those used in scRNA-Seq experiments. Opal IHC, a method of signal amplification that employs incubation with an unconjugated primary, followed by a horseradish peroxidase (HRP)-conjugated secondary, and deposition of Opal fluorophore in the tissue, is an attractive method for detection of very low levels of specific proteins (36). We first tested this method by staining for endogenous levels of the chemokine CXCL9 in the liver sinusoids of mice (Fig. S7A), which showed a signal not readily detected with direct or indirect staining methods. Further, because Opal IHC is well described for the imaging of fixed (FFPE) human tissues (37, 38), we next evaluated whether this method could be expanded upon to achieve multi-parameter imaging of tissues from high containment facilities. Due to the extreme fixation conditions required to inactivate select agents such as the Ebola virus (10% formalin for 8 days), the majority of stainable epitopes are lost in these tissues (39, 40). To overcome this significant technical limitation, we developed the Opal-plex method that is based on the IBEX pipeline. Opal-plex extends the usual fluorophore limitations of Opal by combining multi-plex Opal IHC with cycles of IBEX-based bleaching to eliminate signal from the following LiBH_4_-sensitive dyes (Opal 570, 650, and 690) while utilizing the LiBH_4_-resistant dye (Opal 540) as a fiducial (Fig. 6A, Fig. S7B). Using this approach, we achieved single cell resolution of 10 unique markers in heavily fixed mouse LNs (Fig. 6B, Movie S9).

**Figure 6.**
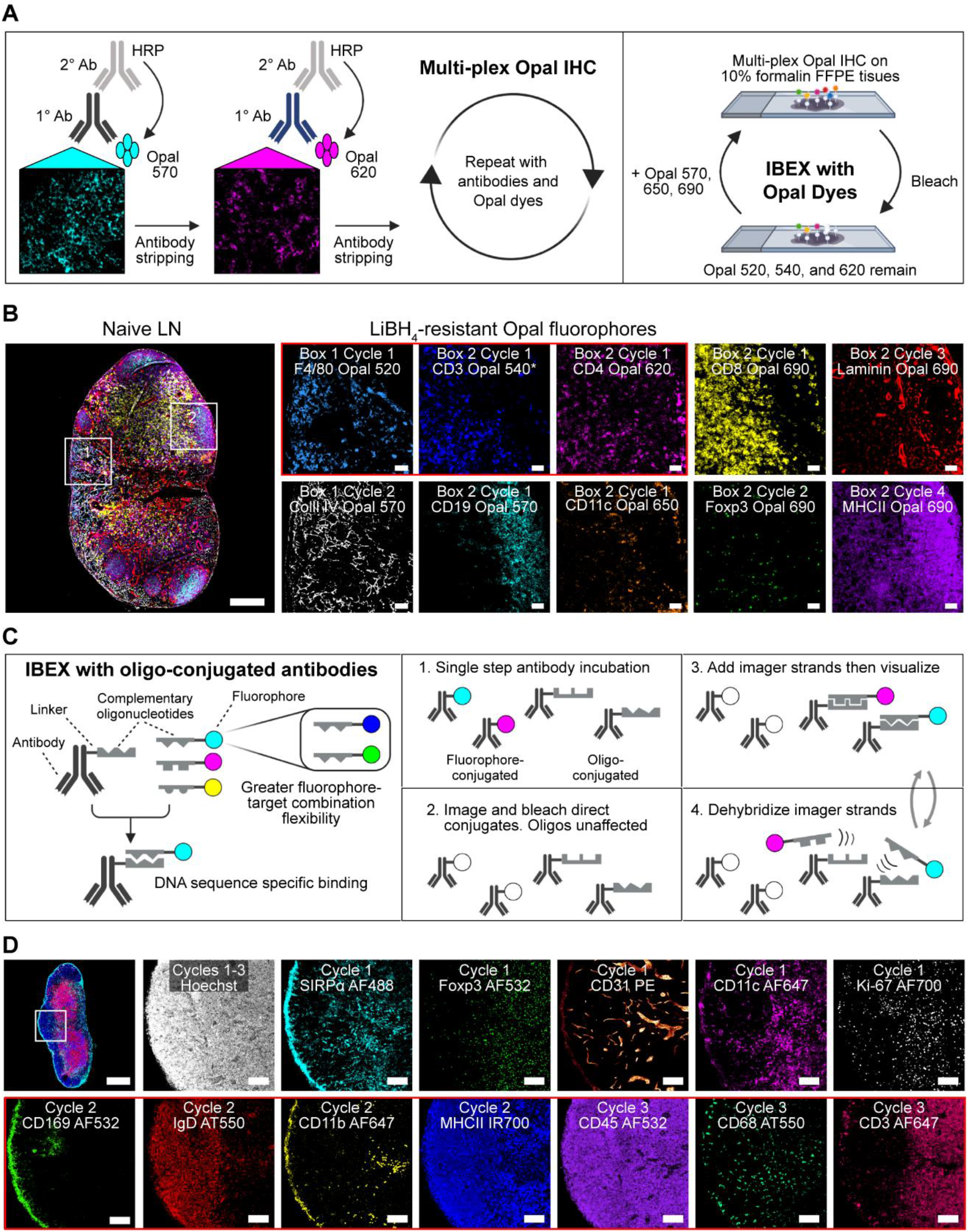
Incorporation of Opal fluorophores and oligo-conjugated antibodies into IBEX workflow. *(A)* Opal-plex imaging method consisting of several rounds of labeling with marker-specific primary antibodies, an HRP-conjugated secondary antibody, Opal dyes, and antibody stripping for each marker-Opal fluorophore pair followed by cycles of IBEX (imaging, removal of coverslip, and bleaching). *(B)* Representative images from a 10 parameter 4 cycle Opal-plex experiment performed on 5 μm FFPE tissue sections from heavily fixed mouse pLNs. CD3 Opal 540 was present throughout cycles 1-4 and served as a fiducial (*). Scale bars (200 μm, left-most panel or 50 μm). (*C)* Schematic depicting principle behind tissue imaging with oligo-conjugated antibodies and incorporation of these reagents into IBEX workflow. (*D)* Confocal images from a 13 parameter 3 cycle IBEX experiment performed on 20 μm tissue sections from an immunized inguinal mouse LN. Cycle 1: Fluorophore-conjugated antibodies. Cycles 2-3: Oligo-conjugated antibodies, Atto550 (AT550). Scale bars (400 μm, top-left panel or 50 μm). Data are representative of 3 similar experiments. See Fig. S7 and Movie S9.

We next evaluated whether oligonucleotide-conjugated antibodies, including those used for CITE-Seq, are compatible with our IBEX workflow. While immunolabeling with oligonucleotide-conjugated antibodies is well established (9), the use of a large number of commercially available TotalSeqA™ antibodies with publicly available oligo-tag sequences, the employment of non-proprietary buffers for hybridization and dehybridization, and the use of a wide spectrum of fluorophore-labeled complementary oligonucleotides provides a truly ‘open source’ system with many advantages. In particular, the imaging method described here applies the same antibodies used for scRNA-Seq, permitting direct comparison between imaging and CITE-Seq datasets while providing a much-needed spatial context for the cell populations identified. Using this approach, we were able to achieve high quality tissue staining with 5 unique fluorophores (Fig. S7C). This method can be directly integrated into our IBEX protocol, alongside fluorophore-conjugated antibodies when CITE-Seq antibodies to desired targets do not exist, as LiBH_4_ bleaching leaves oligonucleotide binding intact (Fig. 6C-D). Importantly, the quality of staining achieved with oligonucleotide-conjugated antibodies, even after multiple cycles of LiBH_4_ bleaching, is comparable to conventional IF as quantitative differences, e.g., higher expression of MHCII on DCs versus B cells, can still be observed (Fig. 6D, Fig. S7D, Movie S9). In summary, this protocol improves upon existing high dimensional DNA-based imaging techniques by offering full flexibility in antibody-fluorophore pairing, integrating commercially produced CITE-Seq reagents, reducing antibody labeling to one step, and extending the number of fluorophores per cycle.

## Discussion

Multi-plex imaging of tissues is increasingly important for studies of tumor-immune interactions, for discovery efforts such as the Human Cell Atlas, for better understanding pathological events in infected or physically damaged tissues, and for placing data from isolated cells in the context of *in situ* tissue organization. IBEX is a broadly applicable technique that utilizes conventional microscopes and commercially available antibodies to obtain these essential high dimensional imaging data. IBEX improves upon existing iterative methods by addressing many of the limitations inherent to these techniques. First, we have significantly reduced the fluorophore inactivation step and antibody labeling time from >16 hours to <1 hour using a rapid chemical bleaching agent and antibody labeling employing a commercial, non-heating microwave. Second, our selection of the bleaching agent LiBH_4_ provides an efficient means to bleach over 15 unique fluorophores while preserving select fluorophores to serve as repeated markers for registration. Importantly, LiBH_4_ treatment does not cause tissue or epitope loss as evidenced by our ability to obtain highly multi-plexed data over several cycles in a wide range of tissues with a very large number of antibodies. Third, and integral to the preservation of tissue integrity through multiple fluid handling cycles, was the use of the tissue adhesive chrome gelatin alum. Importantly, this adhesive adheres delicate tissues to the slide or coverslip surface while maintaining key anatomical features. Finally, the SimpleITK workflow described here represents a significant advancement for the registration of images obtained via cyclic IF methods. In addition to offering flexibility in terms of the repeated markers (membrane, nuclear, structural) used, it provides alignment of markers present on the same cell but not utilized as the fiducial. This is a critical standard for all high dimensional imaging methods because multiple markers are often required to phenotype a particular cell type and staining for the relevant epitopes may occur in different imaging cycles.

In addition to developing an efficient method for highly multi-plexed imaging, the IBEX workflow, unlike commercial all-in-one systems (9–11), offers flexibility in terms of cellular markers, antibody-fluorophore combinations, and microscope configurations employed. Because the chemistry of bleaching depends on the fluorophore and not the antibody to which it is conjugated, once the bleaching conditions are defined, staining panels can be designed using specific combinations of fluorophores without regard for the target epitopes of the antibodies employed, providing the user with extreme versatility in experimental design. To this end, we report the validation of more than 200 commercially available antibodies conjugated to fluorophores with excitation and emission spectra ranging from 405 nm to 750 nm. Furthermore, we demonstrate that commercially available oligonucleotide-conjugated antibodies can be seamlessly integrated into our IBEX workflow, representing the first application of TotalSeqA™ antibodies for *in situ* IHC. Given that the barcode sequences for TotalSeqA™ antibodies are disclosed, and a wide range of fluorophore-conjugated oligos is readily available, fluorophore and antibody pairing can be fully customized to match microscope configuration, epitope abundance, and unique tissue characteristics. Taken together, the oligonucleotide-staining method described here provides a completely ‘open’ method to achieve highly multi-plexed IF imaging using the same antibodies employed for flow cytometry and/or CITE-Seq, enabling effective cross-referencing of datasets derived from these complementary technologies.

As a proof-of-concept, we have used the IBEX workflow to examine such issues as the visualization of difficult to extract myeloid populations in various tissues, changes in immune cell composition following immune perturbation, and detection of low abundance epitopes. For the first application, we were able to visualize tissue-resident macrophages that are difficult to characterize using other methods such as flow cytometry because of their limited recovery upon enzymatic tissue digestion (12). Using the panels of antibodies outlined here, we were able to deeply phenotype medullary (CD169^+^F4/80^+^CD11b^+^Lyve-1^+/−^) and subcapsular sinus (CD169^+^F4/80^−^CD11b^+^) macrophages in the LNs (41) as well as alveolar (SiglecF^+^CD11b^−^CD11c^+^) and interstitial (CD11b^+^CD11c^+^MHCII^+^) macrophages of the lung (42). Additionally, we show that the IBEX method can be scaled to capture ultra-high content imaging in human tissues. The ability to survey large areas of human tissue is critically important as all possible information needs to be extracted to provide maximally useful clinical and research data.

Beyond simple visualization of diverse cell types, we have shown compatibility between IBEX-generated data and downstream single-cell, spatially-resolved analysis using the open source method histoCAT. From the 10 cycle IBEX experiments described above, the histoCAT workflow identified 29 phenotype clusters characterized by the expression of several different markers present in distinct imaging cycles. Importantly, this approach identified well described changes following immunization such as an increase in Tfh cells and GC B cells (43). Finally, incorporation of multi-plex Opal IHC into our IBEX workflow facilitated the detection of low abundance epitopes present in conventionally fixed tissues while aiding in the detection of epitopes lost under extreme fixation conditions. The latter represents a significant achievement because few studies have visualized the immunopathology induced by select agents and, to date, the largest number of parameters examined in a single section has been limited to 3 (44). The ability to use IBEX in its native format and with the Opal-plex variation is especially valuable in the context of the current COVID-19 pandemic. Preliminary data show that these methods work well with highly fixed post-mortem samples from such patients.

In summary, IBEX constitutes a versatile technique for obtaining high content imaging data using conventional microscopes and commercially available antibodies. In addition to providing a valuable resource for studying tissue-based immunity in animal models of disease, ongoing studies have shown the value of the IBEX method to provide a spatially-defined assessment of complex cell phenotypes from diverse organs including lung, kidney, heart, and lymphoid tissues from surgical specimens as well as post-mortem samples from human COVID-19 patients. We believe that the open nature of the reagents that can be utilized, and the variety of instruments suitable for implementation of IBEX, make it an attractive method for many laboratories seeking to obtain a deeper understanding of cell composition and spatial organization in tissues of interest.

## Materials and Methods

Detailed descriptions of animals, immunization and tissue preparations, reagents, imaging protocols with Opal fluorophores and oligonucleotide-conjugated antibodies, microscopy configurations, and image acquisition and analysis details are reported in the SI Materials and Methods.

### Mouse and human tissues

Murine organs and human LNs (1 cm^3^ or smaller in size) were fixed with BD CytoFix/CytoPerm (BD Biosciences) diluted in PBS (1:4) for 2 days. Following fixation, all tissues were washed briefly (5 minutes per wash) in PBS and incubated in 30% sucrose for 2 days before embedding in OCT compound (Tissue-Tek). All mice were maintained in specific pathogen-free conditions at an Association for Assessment and Accreditation of Laboratory Animal Care-accredited animal facility at the National Institute of Allergy and Infectious Diseases (NIAID). All procedures were approved by the NIAID Animal Care and Use Committee (NIH). De-identified human LN samples were obtained from patients undergoing elective risk-reducing gastrectomies or colon resections for colon adenocarcinoma at the National Cancer Institute (NCI) based on an Institutional Review Board (IRB) approved tissue collection protocol (#13C-0076).

### IBEX using inverted confocal microscope

20-30 μm sections were cut on a CM1950 cryostat (Leica) and adhered to 2 well Chambered Coverglasses (Lab-tek) coated with 15 μl of chrome alum gelatin (Newcomer Supply) per well. Frozen sections were permeabilized, blocked, and stained in PBS containing 0.3% Triton X-100 (Sigma-Aldrich), 1% bovine serum albumin (Sigma-Aldrich), and 1% mouse or human Fc block (BD Biosciences). For conventional IF, sections were first blocked 1-2 hours at room temperature and then stained for 12 hours at 4 ° C in a humidity chamber. For microwave-assisted IF, we utilized the PELCO BioWave Pro 36500-230 microwave equipped with a PELCO SteadyTemp Pro 50062 Thermoelectric Recirculating Chiller (Ted Pella). A 2-1-2-1-2-1-2-1-2 program was used for immunolabeling where ‘2’ denotes 2 minutes at 100 watts and ‘1’ denotes 1 minute at 0 watts. The above program was executed once for blocking and secondary antibody labeling and twice for primary antibody labeling. A complete list of antibodies and tissue-specific panels can be found in Tables S1-S5. Cell nuclei were visualized with JOJO-1 (Thermo Fisher Scientific) or Hoechst (Biotium) and sections were mounted using Fluoromount G (Southern Biotech). Mounting media was thoroughly removed by washing with PBS after image acquisition and before chemical bleaching of fluorophores. Samples were treated with 1 mg/mL of LiBH_4_ (STREM Chemicals) prepared in diH_2_O for 15 minutes to bleach all fluorophores except JOJO-1, Hoechst, eF615, and Alexa Fluor 594. To bleach antibodies conjugated to Brilliant Violet 421 (BV421) and Brilliant Violet 510 (BV510) dyes, tissue sections were illuminated using the metal halide lamp with the DAPI filter of the Leica TCS SP8 X inverted confocal microscope. The efficiency of fluorophore bleaching was assessed in real time by viewing the LiBH_4_-incubated samples on the microscope. Following efficient bleaching, the LiBH_4_ solution was removed and samples were washed in 3 exchanges of PBS, restained with the next panel, and mounted with Fluoromount G. Fluorophore inactivation with H_2_O_2_ was conducted as described previously (5) with tissue sections being treated for 1 hour at room temperature with 4.5% H_2_O_2_ (Sigma) prepared in an alkaline solution. Tissue sections were imaged as described in the SI Materials and Methods.

### Image alignment and registration

The alignment of all IBEX panels to a common coordinate system was performed using SimpleITK (23, 24). To facilitate registration, we utilized a common channel present in all panels. As the images may differ by a significant translational motion, we use a Fourier domain-based initialization approach (45) that accommodates for this motion. In addition, we utilized SimpleITK’s multi-scale registration framework with four levels, reducing the resolution by a factor of two per level. Directly using the original voxel sizes, on the order of 10^−3^ mm, in the gradient descent optimizer computations leads to numerical instability. We therefore normalized the voxel dimensions during optimization, while preserving anisotropy. The final, optimal transformations are then used to resample all channels from each panel to the common coordinate system. The software repository for SITK_IBEX can be found on github.com/niaid/sitk-ibex.

### Extensions of IBEX protocol

Integrating IBEX with Opal dyes or oligonucleotide-conjugated antibodies was performed as detailed in the SI Materials and Methods. For multi-plex Opal IHC the following steps—primary antibody incubation, incubation with HRP-conjugated secondary, labeling with Opal dye, and antibody stripping—were repeated to achieve 6-plex imaging using the Opal 520, 540, 570, 620, 650, and 690 fluorophores. After representative images were captured, coverslips were removed and tissue sections were treated with 1 mg/mL of LiBH_4_ prepared in diH_2_O for 30 minutes to bleach the Opal 570, 650, and 690 dyes. Cycles of multi-plex Opal IHC and IBEX were repeated to achieve the desired number of markers. For integration of oligonucleotide-conjugated antibodies into the IBEX workflow, TotalSeq-A™ antibodies were co-incubated with fluorophore-conjugated antibodies which were imaged first before LiBH_4_ bleaching. Oligonucleotides are preserved and complementary fluorophore-conjugated oligonucleotides are used to reveal immunostaining, then either dehybridized or bleached again *in situ* across multiple cycles.

## Acknowledgments

This research was supported by the Intramural Research Program of the NIH, NIAID and NCI. The authors declare no financial conflicts of interest. This research was also partially supported by a Research Collaboration Agreement (RCA) between NIAID and BioLegend, Inc. (RCA# 2020-0333). CJC is supported as a UK-US Fulbright Scholar. We would like to thank Dr. Jagan Muppidi for assistance with SRBC immunization and tissue harvest. Figures were created with Biorender.com.

