## Supplementary Information for "IBEX – A versatile multi-plex optical imaging approach for deep phenotyping and spatial analysis of cells in complex tissues"

[11]Co-first authors

**This PDF includes:**

Supplemental Methods

Figures S1-S7 and Legends

Tables S1-S5

Movies S1-S9 Legends

SI References

### Supplemental Methods

#### Animals, immunizations, and tissue preparations

5-12 week old naïve C57BL/6 and *Cxcl9*$^{-/-}$ mice were purchased from Jackson Laboratories (Bar Harbor, ME) and maintained at a facility at the NIH. Fluorescent Confetti animals were generated by crossing B6.129P2-Gt(ROSA)26Sortm1(CAG-Brainbow2.1)Cle/J × B6.Cg-Tg(UBC-cre/ERT2)1Ejb/2J and LysM-tdTomato mice were generated by crossing LysM-Cre x B6.Cg-Gt(ROSA)26Sortm14(CAG-tdTomato)Hze/J (Jackson Laboratory). Fluorescent Confetti animals heterozygous for both transgenes were injected intraperitoneally (i.p.) with tamoxifen 100 μg per gram of body weight in peanut oil (Sigma-Aldrich) on day 0 and day 2 and tissues were collected on day 4 for processing. To evaluate changes in the immune cell composition and LN architecture following immunization, sheep red blood cells (SRBCs, Colorado Serum Company) were prepared by resuspending 3 mL of SRBCs in 10 mL of Hanks' Balanced Salt Solution (HBSS, Sigma-Aldrich). Mice were injected subcutaneously (s.c.) with SRBCs in a volume of 25-50 μl per site on day 0 and day 4 with organs harvested on day 11. Unless stated otherwise, all tissues were treated similarly and embedded as whole organs. Prior to fixation, murine livers and lungs were perfused with PBS in order to remove blood. The ileal portion of the small intestine was excised and prepared using the Swiss roll technique (1). Livers were collected from LysM-tdTomato mice and processed as described previously (2). Lungs were inflated with 10 ml of fixative via a tracheal cannula before harvest (3). Lungs were then tethered to a small weight and fixed overnight in BD CytoFix/CytoPerm (BD Biosciences) diluted in PBS (1:4). Following fixation, all tissues were washed briefly (5 minutes per wash) in PBS and incubated in 30% sucrose for 2 days before embedding in OCT compound (Tissue-Tek).

#### Additional considerations for IBEX method

For optimal results, it is important to be aware of the expiration date of the chrome gelatin alum and to follow these steps for coating chambers and glass slides: 1) spread the adhesive evenly over the imaging substrate, 2) dry 1 hour at 60° C, 3) section tissue onto coated chamber or glass slide, 4) dry for 1 hour at 37° C, and 5) work with freshly prepared samples (don't store at -20° C). In order to achieve efficient bleaching, always use 1 mg/ml solutions of $LiBH_4$ within 4 hours of preparation and wait until large bubbles form before adding to tissue (usually 10 minutes after dissolving in $diH_2O$). Be careful as $LiBH_4$ can react violently with water. To avoid flames, always work with small amounts (<10 mg) of $LiBH_4$. Store $LiBH_4$ with desiccant and use a new vial of $LiBH_4$ after 4 weeks of use. As noted in the "IBEX using inverted confocal microscope" section of the Methods section, Brilliant Violet 421 (BV421) and Brilliant Violet 510 (BV510) dyes require $LiBH_4$ bleaching in the presence of light (metal halide lamp with the DAPI filter). To ensure efficient bleaching of BV421 and BV510 conjugates, we assess bleaching in real time by viewing the $LiBH_4$-incubated samples under the microscope, bleaching a field of view (1-2 minutes), and moving to an adjacent field of view (1-2 minutes). This process is repeated until the entire region of interest is bleached. Of note, the time required for this process is dependent on the imaging area and objective used for bleaching with larger areas requiring longer overall bleaching times. As an example, a tissue section from a mouse LN (1-2 mm) can be readily bleached in 15 minutes using this approach. Another consideration is that all images be oriented identically in the Z stack (same begin and end) for proper alignment with SimpleITK. For gross observation and visualization, we found that IBEX-generated images could be aligned using Imaris 9.5.0 (Bitplane) by "adding images" and manually aligning images with the reference manager. Alternatively, the image alignment feature under the image processing menu was used to provide a rough assessment of image registration before submitting IBEX-generated images to the SimpleITK workflow.

**Ensuring optimal imaging quality: Antibody validation and controls**

Selection of antibodies that bind their targets with great specificity while yielding reproducible immunolabeling across different samples is critical for all imaging methods. Wherever possible, we utilize highly cited antibodies previously validated for immunofluorescence of fixed frozen tissues. Upon identifying suitable antibody candidates for our imaging workflow, we note the location (membrane, cytoplasm, nucleus) and tissue distribution of the marker of interest. Additionally, we procure positive and negative control tissues based on the described expression of a particular marker. Finally, we pair the new antibody with previously validated markers that co-stain the same cell type. For example, the SPARC antibody (R&D AF941) is reported to label macrophages, fibroblasts, and endothelial cells within human tissues. To validate this antibody, we evaluate the spatial distribution of the anti-SPARC antibody in human LNs co-stained with CD3 (negative) and CD11c and CD31 (positive controls). Prior to all iterative imaging, antibodies are tested for their sensitivity to $LiBH_4$ by pretreating tissue sections with $LiBH_4$ for 15 minutes. To evaluate whether epitopes are sensitive to $LiBH_4$, staining patterns for individual antibodies are compared between serial sections with or without $LiBH_4$ pre-treatment. As an additional control, individual panels are acquired on serial sections in parallel with iterative rounds of imaging. The spatial distribution patterns are then compared between the serially and iteratively acquired images for each antibody to ensure there is no epitope loss or steric hindrance with cyclic imaging. Finally, antibody concentrations vary greatly depending on the tissue, fixation conditions, and imaging system employed. For these reasons, we strongly recommend careful titration of all antibodies prior to IBEX imaging.

**IBEX imaging conditions for inverted confocal microscope**

Representative sections from different tissues were acquired using an inverted Leica TCS SP8 X confocal microscope equipped with a 40X objective (NA 1.3), 4 HyD and 1 PMT detectors, a white light laser that produces a continuous spectral output between 470 and 670 nm as well as 405, 685, and 730 nm lasers. Panels consisted of antibodies conjugated to the following fluorophores and dyes: Hoechst, BV421, BV510, AF488, AF532, JOJO-1, PE, eF570, AF555, AF594, AF647, eF660, and AF700. All images were captured at an 8-bit depth, with a line average of 3, and 1024x1024 format with the following pixel dimensions: x (0.284 μm), y (0.284 μm), and z (1-1.25 μm). Images were tiled and merged using the LAS X Navigator software (LAS X 3.5.5.19976). To ensure proper alignment over distinct imaging cycles, careful attention was paid to the quality of image stitching achieved with the Leica software and z-stacks were set by manual inspection of notable features such as unusually shaped nuclei throughout the tissue volume. These unusual features were matched across the z-stack and over multiple cycles of IBEX.

**Adoption of IBEX to additional imaging systems**

For adoption of the IBEX protocol to an upright Leica TCS SP8 X confocal microscope, 20-30 μm sections were adhered to Super Frost Plus Gold slides (Electron Microscopy Services) coated with 30 μl of chrome alum gelatin. The IBEX protocol was executed as described above with the following exceptions: slides were mounted with a No. 1.5 coverslip (VWR) and antibody panels were designed without BV421 and BV510 conjugated antibodies. These conjugates were omitted because they require the tissue to be immersed in $LiBH_4$ while illuminated with the metal halide lamp, an impossibility for samples mounted on slides. Following image acquisition, coverslipped slides were immersed in PBS until the coverslip floated off. Non-coverslipped slides were incubated with $LiBH_4$ for 15 minutes, washed extensively in PBS, immunolabeled with the next round of antibodies, mounted with Fluoromount G, and coverslipped. Image acquisition parameters and system configurations were identical to the details listed for the inverted Leica TCS SP8 X confocal microscope with the exception of the 685 nm laser. For adoption of the IBEX protocol to an inverted fluorescent microscope, 5-10 μm sections were adhered to Super Frost Plus Gold slides coated with chrome alum gelatin. Slides were mounted with a coverslip and antibody panels were designed with the following fluorophores and dyes: Hoechst (Biotium), AF488, PE, and AF647. Representative sections were acquired using a Keyence BZ-X800 microscope equipped with a 40X objective, metal halide lamp, and DAPI, GFP, TRITC, and Cy5 filter sets. All images were captured using the auto-exposure settings for each channel at standard resolution yielding an 8-bit image. Haze reduction was applied post-acquisition to improve image quality.

**Image analysis and quantification**

Fluorophore emission was collected on separate detectors with sequential laser excitation of compatible fluorophores (3-4 per sequential) used to minimize spectral spillover. The Channel Dye Separation module within the LAS X 3.5.5.19976 (Leica) was then used to correct for any residual spillover. Threshold identification, voxel gating, surface creation, and masking were performed as previously described (4, 5). For publication quality images, gaussian filters, brightness/contrast adjustments, and channel masks were applied uniformly to all images. Unless stated otherwise, images are presented as maximum intensity projections (MIP) of tiled z-stacks. Quantification of cell surfaces was based on images with unadjusted gamma values. To calculate the amount of signal remaining after $LiBH_4$ bleaching (Figs. 1D and S7B), the Color Pixel Counter plugin developed by Ben Pichette for FIJI was used (6). The Structural SSIMilarity (SSIM) index, a method for measuring the similarity between 2 images, was used to assess whether $LiBH_4$ treatment removed primary antibodies from the tissues (7) (Fig. S2). The surface creation module of Imaris 9.5.0 (Bitplane) was used to segment cells based on the nuclear marker Hoechst (Figs. S1C or S5B) or CFP+, GFP+, YFP+, or RFP+ expression (Fig. S4B). For consistency, segmentation conditions were applied as a batch using identical parameters. Segmented cells were randomly colored and manually inspected to assess the quality of segmentation based on Hoechst or FP staining. For Fig. S5B, segmentation on Hoechst+ cells provided accurate identification of round, regular shaped, lymphocytes but, as expected, struggled with morphologically complex stromal and structural cell segmentation. Nevertheless, this approach provided a relative quantification for the number of surfaces positive for each individual marker. Importantly, these qualitative measures were in agreement with the amount of signal present by visual inspection. Surfaces were scored positive for a given marker based on absolute intensity values generated in Imaris and these values were applied uniformly to serial and IBEX-generated images to quantify marker-positive surfaces (Fig. S5B).

**histoCAT analysis of IBEX-generated images**

Using the workflow described previously (8), IBEX-generated images were segmented using Ilastik (Version 1.3.3) (9) and CellProfiler (Version 3.1.9) (10). Briefly, B220, CD3, and CD45 (most of the immune cells) markers were merged into a single membrane and JOJO-1 was used as a nuclear marker. Membrane and nuclear Images were loaded into Ilastik for supervised training of pixel segmentation. The trained Ilastik model classified image pixels into 3 classes (membrane, nuclear, and background) and generated probability maps for CellProfiler for cellular objects segmentation. CellProfiller-generated object/mask images were exported as TIFF files for downstream analysis. To quantify marker expression, individual marker images, along with masks, were loaded into histoCAT and Phenograph clustering with default parameters was performed. Phenograph consistently identified 29 different clusters/phenotypes (Fig. 4B). Hierarchical clustering and heatmap (Fig. S6B) were generated from the Phenograph output using the Seaborn

Python package. The high-dimensional single-cell data was projected onto 2 dimensions using the t-SNE module in histoCAT for visualization purposes shown in Fig. 4B. Cellular populations identified in Fig. 4C were manually phenotyped based on the cellular markers expressed within each Phenograph cluster and summarized in the heat map found in Fig. S6B: $CD4^+$ T (cluster 3), $CD8^+$ T (cluster 2), Tfh (cluster 19), Tregs (cluster 18), naïve B (cluster 5), $MHCII^{Hi}$ B (cluster 13), PCs (cluster 10), GC B (cluster 6), $CD68^+$ macrophages (cluster 15), $CD206^+$ macrophages (cluster 11), SCS macrophages (cluster 17), MS macrophages (cluster 25), pan DC (cluster 4), cDC1/dDCs (cluster 22), and cDC2s (cluster 26). A median of ratios method (11) was performed for the cell count normalization (Fig. S6D). All histoCAT analysis was done on an iMAC, (Retina 5K, 27-inch, 2017, 4.2Ghz i7, 64G RAM) with macOS Mojave operating system.

**Multi-plex Opal IHC on heavily fixed tissues**

pLNs were collected from naïve mice and fixed at 4° C for 8 days in 10% neutral-buffered formalin (Cancer Diagnostics). Following fixation, samples were washed in PBS to remove formalin and embedded in paraffin blocks. 5 μm sections were cut from paraffin blocks, mounted on slides, and left at 37° C for 16 hours to dry. Slides were incubated at 60° C for 45 minutes, dewaxed using a standard protocol of 10 minutes in xylene (2 times), and rehydrated with graded concentrations of ethanol and water (100% ethanol for 10 minutes, 95% ethanol for 10 minutes, 70% ethanol for 5 minutes). Following dewaxing, slides were first rinsed in water and tissues were fixed to the slides by placing in 10% formalin for 15 minutes, rinsed in water, and then placed in TBST (1X TBS + 0.5% Tween20 (Thermo)) to prevent drying out. Antigen retrieval was performed using the AR6 buffer from the Opal Multi-plex kit (Akoya Biosciences) using a conventional microwave at 100% power for 45 seconds followed by 10% power for 15 minutes. To perform multi-plex Opal IHC, tissue sections were first blocked in Opal antibody diluent/blocking buffer (Akoya Biosciences) for 10 minutes. Next, unconjugated primary antibodies were added to the tissue for 12-16 hours at 4° C, 4 hours at room temperature, or 30 minutes using the PELCO BioWave Pro 36500-230 microwave described under 'IBEX using inverted confocal microscope' section of the Material and Methods. After primary antibody incubation, samples were washed with TBST and an Opal anti-rabbit HRP-conjugated secondary antibody was added at a dilution of 1:5 for 10 minutes at room temperature or with a species matched HRP-conjugated secondary antibody for 1 hour at room temperature. Samples were washed several (5-6) times with TBST and Opal dyes (diluted 1:100 in 1X amplification buffer (Akoya Biosciences)) were added to the sample and incubated for 10 minutes at room temperature. Following this last step, samples were washed with TBST and antigen retrieval/antibody stripping was performed using the AR6 buffer and conventional microwave treatment described above. This series of steps—primary antibody incubation, incubation with HRP-conjugated secondary, labeling with Opal dye, antibody stripping—was repeated for the following Opal fluorophores to achieve 6-plex imaging: Opal 520, 540, 570, 620,

650, and 690. Slides were mounted and imaged with an upright Leica TCS SP8 X confocal microscope as described above. After representative images were captured, coverslips were removed and tissue sections were treated with 1 mg/mL of $LiBH_4$ prepared in $diH_2O$ for 30 minutes to bleach the Opal 570, 650, and 690 dyes. Cycles of multi-plex Opal IHC and IBEX were repeated to achieve the desired number of markers. Individual images were aligned and processed as described above. A complete list of antibodies and reagents can be found in Table S4.

**Chemokine staining for endogenous CXCL9**

Livers were collected from naïve WT and *Cxcl9*$^{-/-}$ animals, perfused with 1% PFA through the portal vein, and fixed at 4° C for 12-16 hours in BD CytoFix/CytoPerm diluted 1:4 in PBS. Samples were incubated in sucrose for 24 hours at 4° C and frozen in OCT. To visualize endogenous CXCL9 levels, 20 µm sections were rehydrated in PBS, incubated in 0.1% $H_2O_2$ for 30 minutes to quench endogenous peroxidase, and incubated with unconjugated (CXCL9) and fluorescently-conjugated primary antibodies. Tissue sections were washed to remove unbound antibodies, fixed with 10% formalin to cross-link the bound antibodies to the tissue, washed with TBST, and an Opal anti-rabbit HRP-conjugated secondary antibody was added at a dilution of 1:5 for 10 minutes at room temperature. Following secondary antibody incubation, samples were washed in TBST and the Opal 650 dye (diluted 1:100 in 1X amplification buffer) was added for 5 minutes. Slides were washed, mounted in Fluoromount G, and imaged as described above. A complete list of antibodies and reagents can be found in Table S4.

**Incorporation of TotalSeqA$^{TM}$ antibodies into IBEX workflow**

LNs from C57BL/6 mice were harvested and processed as described above. Integrating published methods, slides containing 30 µm sections were blocked for 30 minutes at room temperature in Buffer A (PBS supplemented with 5% donkey serum (Jackson Immunoresearch), 200 µg/ml sheared salmon sperm (Thermo), 0.2% Triton X-100 (Sigma) and 5 µg/ml Fc-block (Thermo)). Immunolabeling was conducted using Buffer A with the addition of 5 mM EDTA and 0.02% Dextran sulfate, TotalSeqA$^{TM}$ antibodies (5 µg/ml), and fluorophore-conjugated antibodies at the indicated concentrations (Table S1) (12). All primary antibody incubations were performed using the PELCO BioWave Pro 36500-230 microwave as described in the "IBEX using inverted confocal microscope' section of the Materials and Methods. Following antibody incubation, slides were washed using PBS with 0.1% Triton X-100 and post-fixed with 5 mM of BS(PEG)5 (Thermo) for 10 minutes before quenching with 100 mM Ammonium Chloride (Sigma). After washing with PBS, Fluoromount G was applied and fluorescently-conjugated antibodies were imaged in the first cycle. Mounting medium was removed by washing with PBS before bleaching with $LiBH_4$ for 15 minutes. For *in situ* visualization of TotalSeqA$^{TM}$ antibodies, oligonucleotides were designed complementary to the TotalSeqA™ barcode. These complementary oligonucleotides were synthesized by Integrated DNA Technologies (IDT) and 5'-end conjugated with indicated fluorophores before HPLC purification (Table S5). Direct labeling of TotalSeqA™ antibodies with complementary fluorescent oligonucleotides was achieved by adding 1 µM concentrations of imager oligonucleotides in PBS with 0.1% Triton X-100 and incubating for 30 minutes at room temperature. Tissue sections were washed with 0.5X PBS and 0.1% Triton X-100, mounted, and imaged (Cycle 2 of IBEX). After removal of Fluoromount G, oligonucleotides were dehybridized for 15 minutes by incubation with a 0.1X PBS solution containing 30% Formamide. Following 5-6 exchanges of PBS, the next set of oligonucleotides (Cycle 3) was applied as before.

### Figures S1-S7 and Legends

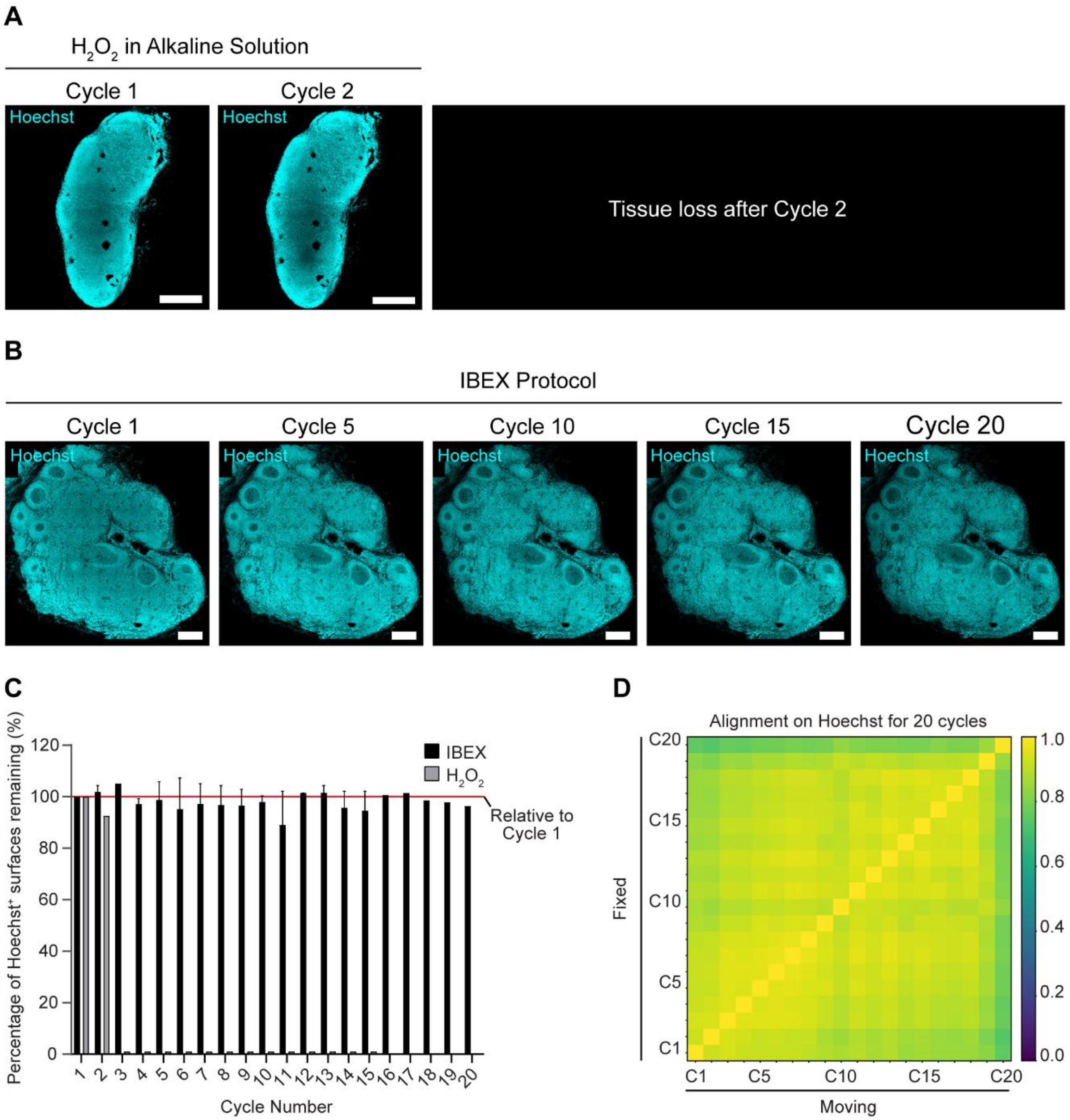


**Figure S1. IBEX protocol preserves tissue integrity over high cycle number.**

*(A)* Confocal images of mouse inguinal LN iteratively imaged using $H_2O_2$ in alkaline solution. Tissues lifted after 2 cycles and could not be imaged as a comparison. Scale bar corresponds to 500 µm. *(B)* Confocal images of human mesenteric LN iteratively imaged using IBEX protocol for 20 cycles. Scale bar corresponds to 500 µm. *(C)* Quantification of percentage of Hoechst+ surfaces remaining per cycle relative to the number of surfaces present in cycle 1. Red line is reflective of no tissue loss. Data are pooled from 2 similar experiments: a 15 cycle mouse inguinal LN and a 20 cycle human mesenteric LN. Shown is the mean ± SEM. *(D)* Cross correlation similarity matrix after alignment with Hoechst across 20 cycle experiment shown in B.

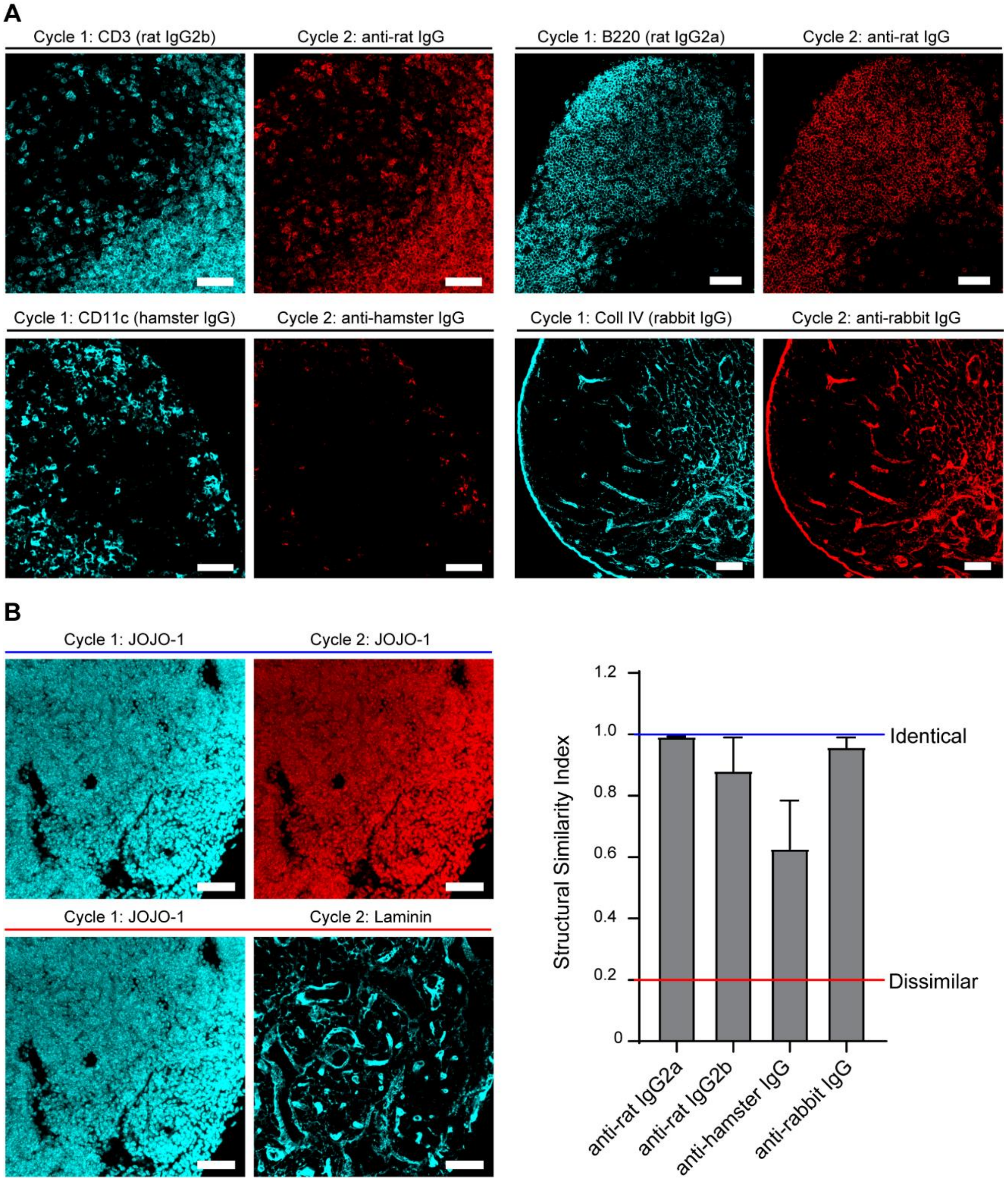


**Figure S2. $LiBH_4$ primarily acts by bleaching fluorophores and not stripping antibodies.**

Popliteal LN sections were stained with primary antibodies directed against various antigens, bleached with $LiBH_4$, and then labeled with secondary antibodies directed against the primary antibody isotype. *(A)* Confocal images of murine LN sections depicting the level of signal present with the primary antibodies as compared to an appropriate secondary antibody after IBEX protocol. *(B)* Bar graph summarizing the level of similarity between images captured before and after $LiBH_4$ treatment. Red line represents dissimilar images (Cycle 1 JOJO-1 versus Cycle 1 Laminin) and blue line denotes highly similar images (Cycle 1 JOJO-1 vs Cycle 2 JOJO-1). Data representative of 5 similar experiments. Shown is the mean ±SEM. Scale bar represents 50 µm.

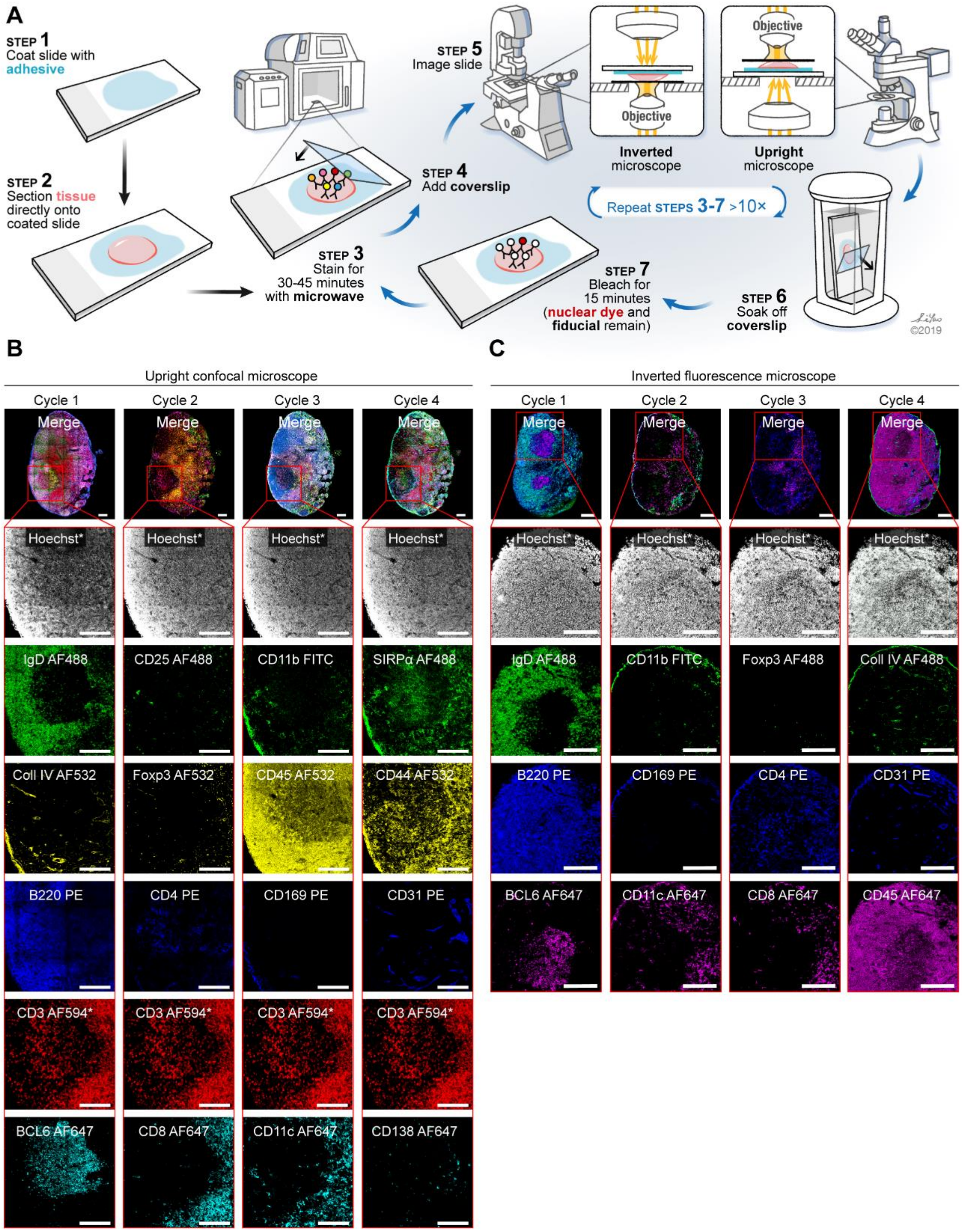


*Fiducials: Hoechst and CD3 AF594, scale bar 150 um

**Figure S3. IBEX can be easily adapted to other microscope configurations.**

*(A)* Schematic depicting IBEX protocol using an inverted or upright microscope. *(B)* Confocal images of popliteal LNs from SRBC-immunized mice. 30 µm tissue sections were labeled with 4 separate 6 parameter imaging panels. The nuclear dye Hoechst and membrane label CD3 AF594 were present throughout cycles 1-4 and served as fiducials. Top panels are a merge of all channels

for each cycle except for Hoechst. Scale bar represents 150 µm. Confocal images were acquired by an upright confocal microscope. (C) 5 µm sections from popliteal LNs from SRBC-immunized mice were visualized using an inverted epifluorescence microscope. Tissue sections were labeled with 4 separate 4 parameter imaging panels with Hoechst serving as a fiducial for cycles 1-4. Scale bar represents 150 µm.

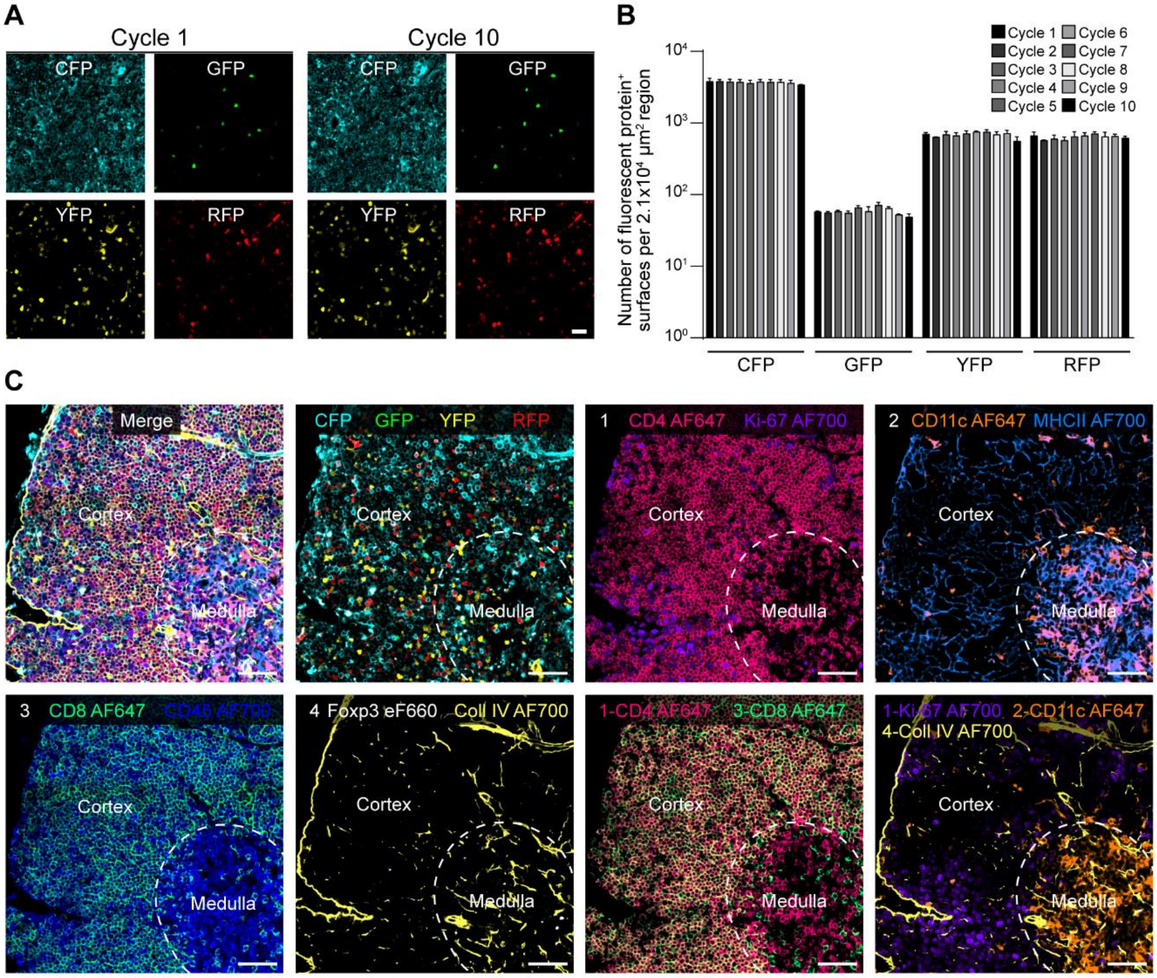


**Figure S4. Fluorescent proteins are not bleached with IBEX protocol.**

Fluorescent Confetti animals were injected i.p. with tamoxifen on day 0 and day 2 to induce the expression of the following fluorescent proteins: membrane CFP, nuclear GFP, and cytoplasmic YFP and RFP. On day 4, tissues were collected and processed for confocal microscopy. *(A)* Example images of signal from fluorescent proteins at Cycle 1 (no $LiBH_4$) and Cycle 10 (after 9 rounds of $LiBH_4$ bleaching). *(B)* Quantification of the number of fluorescent protein+ surfaces per $2.1x10^4$ $\mu m^2$ region of interest from 2 independent experiments. Shown is the mean ± SEM. *(C)* Four cycles of IBEX were applied to 30 µm sections of thymus tissue. AF647 and AF700 conjugated antibodies were used to stain immune and structural markers in the tissue. Representative images showing the compatibility of IBEX with transgenic animals expressing fluorescent proteins. Data are representative of 2 independent experiments. For A and C, images represent a single z slice and scale bar corresponds to 25 µm *(A)* or 50 µm *(B)*.

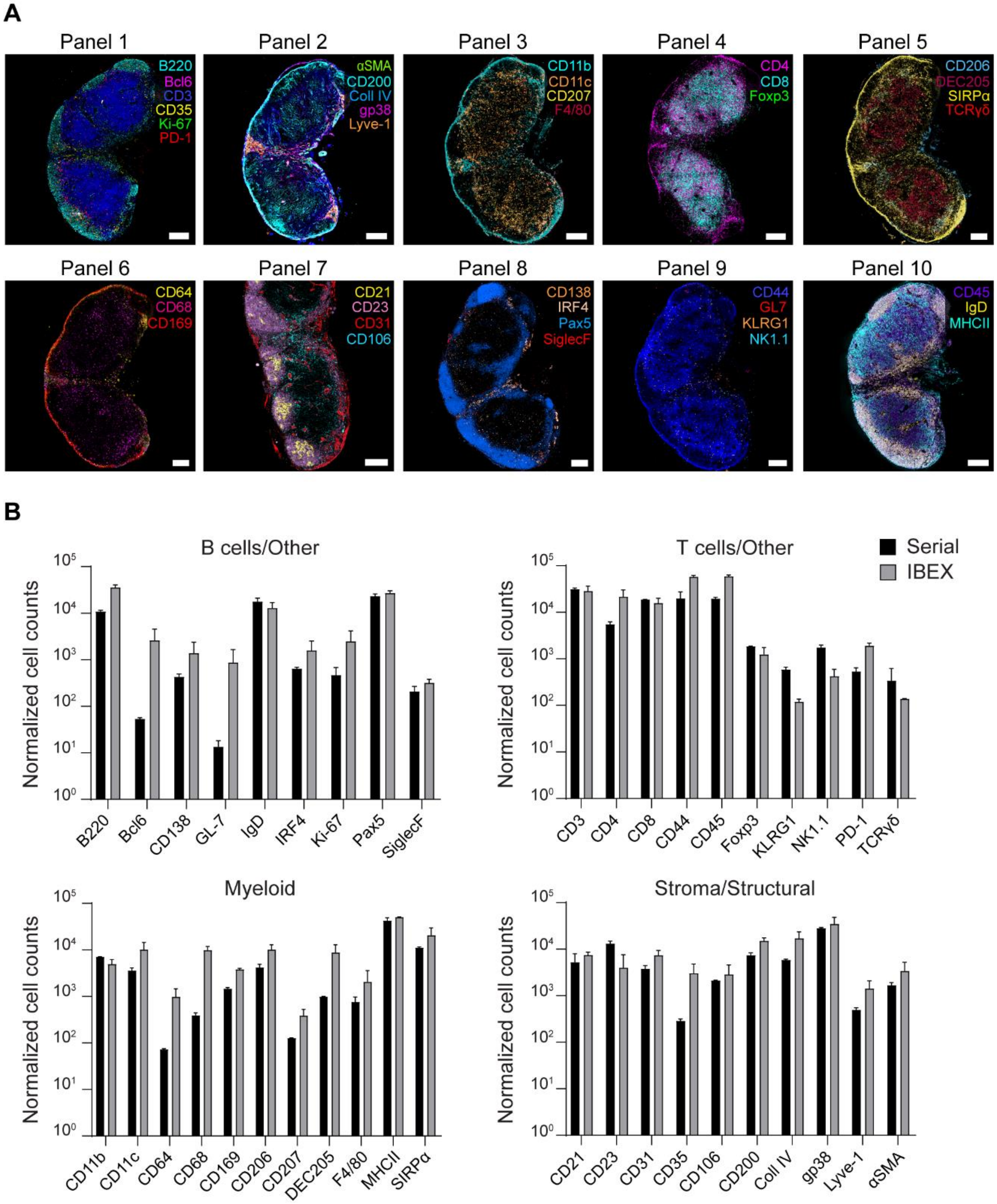


**Figure S5. Comparable staining observed by serial and iterative immunofluorescence methods.**

*(A)* Confocal images of inguinal LNs (iLNs) from SRBC-immunized mice. Serial sections were stained with the same panels used for the 10 cycle 41 parameter IBEX experiments described in Fig. 4A. Scale bar is 200 µm. (*B)* Cells were segmented on the nuclear marker JOJO-1 and the number of surfaces positive for each marker were quantified from images acquired serially as in A or iteratively via IBEX method. Data are from 2-3 LNs per group with 2 immunized iLNs for serial

method and 1 naïve pLN, 1 immunized pLN, and 1 immunized axillary LN (aLN) for IBEX method. Shown is the mean ± SEM. See Movie S7.

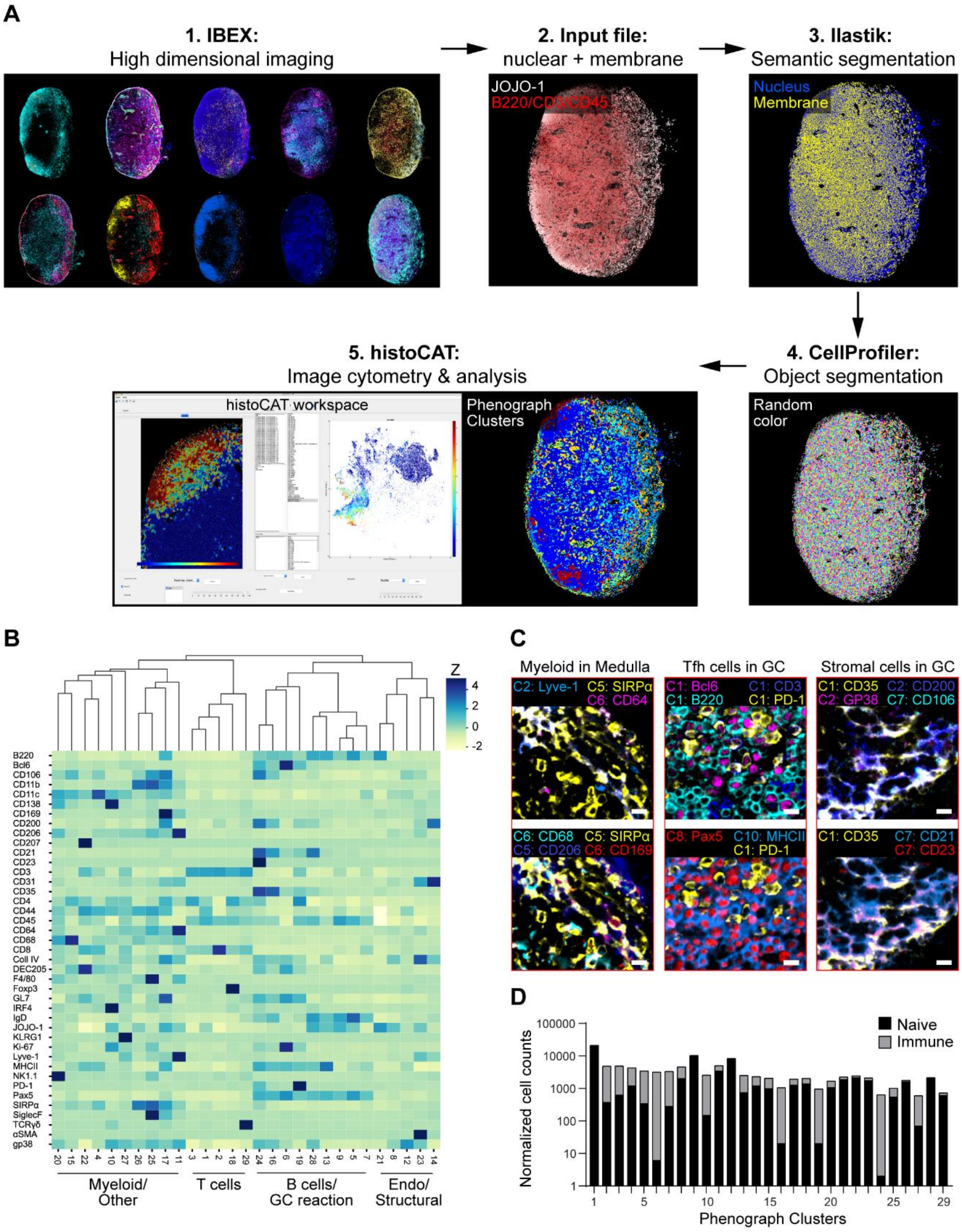


**Figure S6. Data visualization and quantification with histoCAT.**

*(A)* histoCAT workflow for image analysis. Step 1: High dimensional imaging of mouse LNs using IBEX method. Step 2: Nuclei were defined by JOJO-1 and membranes were generated by combining B220, CD3, and CD45 into one composite channel. Step 3: Semantic segmentation was performed using Ilastik. Step 4: Objects were further segmented using CellProfiler. Each

segmented cell is randomly colored and projected back onto original x-y coordinates. Step 5: Data visualization and analysis with histoCAT. (B) Marker expression heatmap for the phenotypes identified by PhenoGraph clustering in histoCAT using segmented cells from naïve (n = 32,091) and SRBC-immunized mouse LNs (n = 80,355). The heatmap displays relative expression levels based on Z-score normalized marker intensity values, and single cells are hierarchically clustered within each phenotype group. Numbers at the bottom of the heatmap indicate corresponding Phenograph cluster IDs with labels corresponding to manually assigned cell populations. Endothelial (Endo). *(C)* Example images of myeloid, Tfh, and stromal cell populations identified by histoCAT Phenograph clustering from an immunized mouse LN. Scale bar corresponds to 10 µm. *(D)* Normalized cell counts for Phenograph clusters obtained from naïve and SRBC-immunized mouse LNs. Data are from one experiment and are representative of 2 similar experiments.

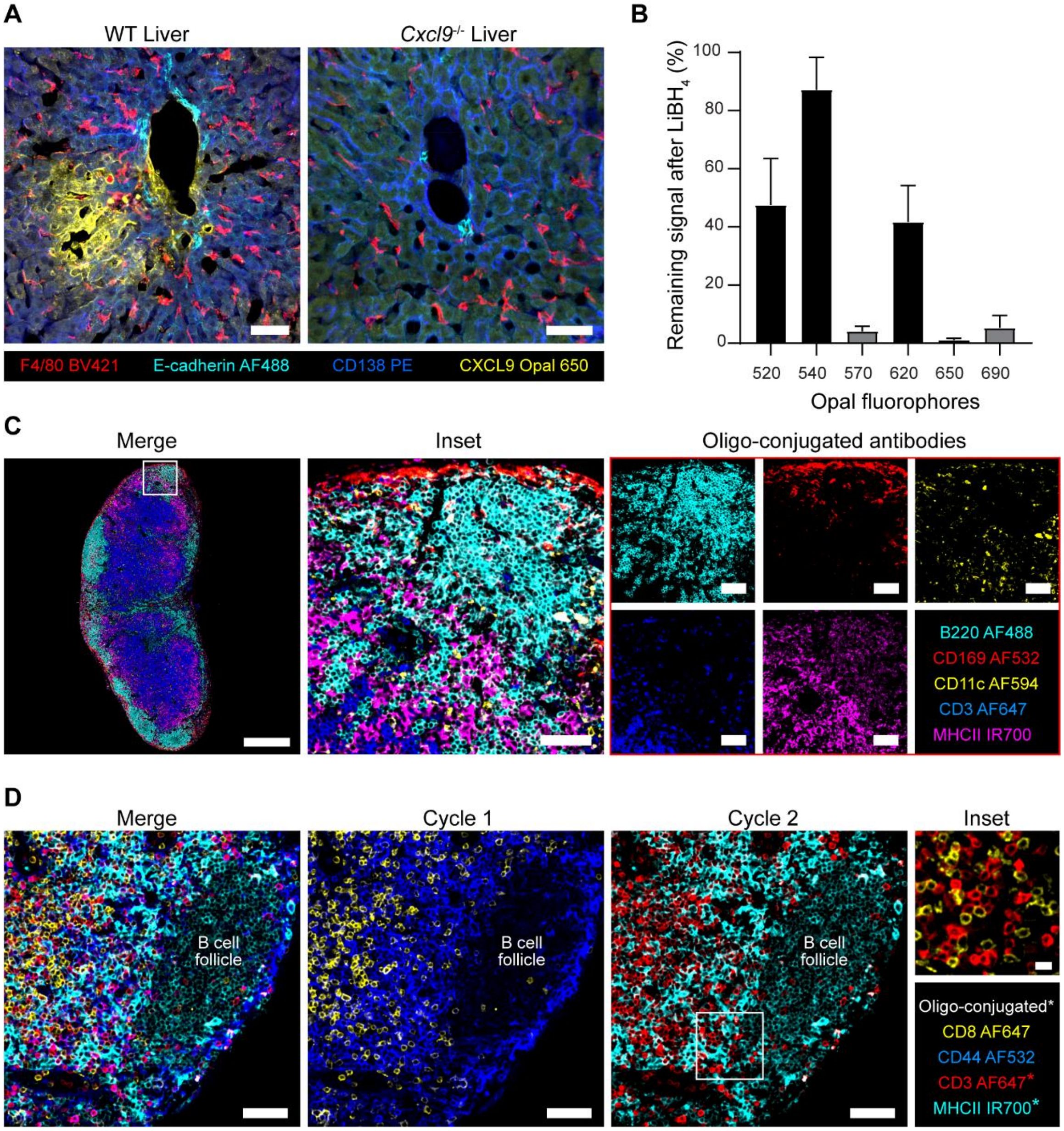


**Figure S7. Extensions of IBEX workflow to enable single cell resolution of endogenous chemokines and immune populations using Opal fluorophores and oligonucleotide-conjugated antibodies.**

*(A)* Detection of endogenous CXCL9 levels in fixed frozen mouse liver sections stained with indicated markers. Scale bars 50 µm. Confocal images are representative of 3 similar experiments. *(B)* Percentage of fluorophore signal remaining after 30 minutes of $LiBH_4$ treatment. Data are pooled from 4 experiments with iterative staining of the multiple Opal fluorophores. Shown is the mean ± SEM. *(C)* Confocal images of mouse inguinal LN labeled with 5 different oligo-conjugated antibodies and complementary fluorescent imager strands. Scale bars (left-most panel 400 µm or 100 µm). (*D)* Representative images from a 2 cycle IBEX experiment performed on inguinal mouse LNs where the first cycle consisted of fluorophore-conjugated antibodies and the second cycle consisted of oligo-conjugated antibodies denoted by the asterisks. Scale bars (50 µm, Inset 10 µm).

### Tables S1-S5

**Table S1. Time and method used to bleach fluorescently conjugated antibodies and dyes.**

| | | | | | | | Time to Bleach (Minutes) | |
|---|---|---|---|---|---|---|---|---|
| **Marker** | **Clone** | **Conjugate** | **Vendor** | **Cat No.** | **Isotype** | **Dilution** | $LiBH_4$ | $LiBH_4$ + Light |
| β-3 Tubulin | AA10 | BV421 | BioLegend | 657412 | Mouse IgG2a, κ | 1:200 | - | <15 |
| β-3 Tubulin | AA10 | AF532 | BioLegend | Custom | Mouse IgG2a, κ | 1:100 | <15 | - |
| B220 | RA3-6B2 | BV421 | BioLegend | 103251 | Rat IgG2a, κ | 1:400 | - | <15 |
| B220 | RA3-6B2 | BV510 | BioLegend | 103248 | Rat IgG2a, κ | 1:300 | - | <15 |
| B220 | RA3-6B2 | AF488 | BD Biosciences | 557669 | Rat IgG2a, κ | 1:400 | <15 | - |
| B220 | RA3-6B2 | AF532 | Thermo | 58-0452-82 | Rat IgG2a, κ | 1:50 | <15 | - |
| B220 | RA3-6B2 | PE | BD Biosciences | 553090 | Rat IgG2a, κ | 1:400 | <15 | - |
| B220 | RA3-6B2 | eF570 | Thermo | 41-0452-80 | Rat IgG2a, κ | 1:200 | <15 | - |
| B220 | RA3-6B2 | eF615 | Thermo | 42-0452-82 | Rat IgG2a, κ | 1:200 | >120 | - |
| B220 | RA3-6B2 | AF647 | BioLegend | 103226 | Rat IgG2a, κ | 1:400 | <15 | - |
| B220 | RA3-6B2 | AF700 | BioLegend | 103232 | Rat IgG2a, κ | 1:50 | <15 | - |
| BCL2 | 100 | AF647 | BioLegend | 658705 | Mouse $IgG_1$ | 1:25 | <15 | - |
| Bcl6 | K112-91 | AF647 | BD Biosciences | 561525 | Mouse $IgG_1$, κ | 1:50 | <15 | - |
| CD1c | L161 | PE | BioLegend | 331506 | Mouse $IgG_1$, κ | 1:50 | <15 | - |
| CD1d | 1B1 | PE | BD Biosciences | 553846 | Rat IgG2b, κ | 1:100 | <15 | - |
| CD3 | 17A2 | BV421 | BioLegend | 100228 | Rat IgG2b, κ | 1:400 | - | <15 |
| CD3 | 17A2 | BV510 | BioLegend | 100234 | Rat IgG2b, κ | 1:50 | - | <15 |
| CD3 | 17A2 | AF488 | BioLegend | 100210 | Rat IgG2b, κ | 1:200 | <15 | - |
| CD3 | 17A2 | AF532 | Thermo | 58-0032-80 | Rat IgG2b, κ | 1:50 | <15 | - |
| CD3 | 17A2 | PE | BioLegend | 100205 | Rat IgG2b, κ | 1:200 | <15 | - |
| CD3 | 17A2 | AF594 | BioLegend | 100240 | Rat IgG2b, κ | 1:400 | >120 | - |
| CD3 | 17A2 | AF647 | BD Biosciences | 557869 | Rat IgG2b, κ | 1:400 | <15 | - |
| CD3 | UCHT1 | AF532 | Thermo | 58-0038-42 | Mouse $IgG_1$, κ | 1:50 | <15 | - |
| CD3 | UCHT1 | AF594 | BioLegend | 300446 | Mouse $IgG_1$ | 1:200 | >120 | - |
| CD4 | GK1.5 | BV421 | BioLegend | 100443 | Rat IgG2b, κ | 1:200 | - | <15 |
| CD4 | GK1.5 | BV510 | BioLegend | 100449 | Rat IgG2b, κ | 1:50 | - | <15 |
| CD4 | GK1.5 | PE | BD Biosciences | 553730 | Rat IgG2b, κ | 1:200 | <15 | - |
| CD4 | GK1.5 | AF594 | BioLegend | 100446 | Rat IgG2b, κ | 1:200 | >120 | - |
| CD4 | RM4-5 | AF488 | BD Biosciences | 557667 | Rat IgG2a, κ | 1:100 | <15 | - |
| CD4 | RM4-5 | AF532 | Thermo | 58-0042-80 | Rat IgG2a, κ | 1:50 | <15 | - |
| CD4 | RM4-5 | eF570 | Thermo | 41-0042-82 | Rat IgG2a, κ | 1:100 | <15 | - |
| CD4 | RPA-T4 | AF532 | Thermo | 58-0049-42 | Mouse $IgG_1$, κ | 1:25 | <15 | - |
| CD4 | RPA-T4 | AF700 | BioLegend | 300526 | Mouse $IgG_1$, κ | 1:25 | <15 | - |
| CD8 | 53-6.7 | BV421 | BioLegend | 100738 | Rat IgG2a, κ | 1:200 | - | <15 |
| CD8 | 53-6.7 | BV510 | BioLegend | 100752 | Rat IgG2a, κ | 1:200 | - | <15 |
| CD8 | 53-6.7 | AF488 | BioLegend | 100723 | Rat IgG2a, κ | 1:200 | <15 | - |
| CD8 | 53-6.7 | PE | BD Biosciences | 553032 | Rat IgG2a, κ | 1:400 | <15 | - |
| CD8 | 53-6.7 | AF594 | BioLegend | 100758 | Rat IgG2a, κ | 1:200 | >120 | - |
| CD8 | 53-6.7 | AF647 | BioLegend | 100724 | Rat IgG2a, κ | 1:200 | <15 | - |
| CD8 | SK1 | AF488 | BioLegend | 344716 | Mouse $IgG_1$, κ | 1:50 | <15 | - |
| CD10 | FR4D11 | PE | Caprico Biotechnologies | 103926 | Mouse $IgG_1$, κ | 1:50 | <15 | - |
| CD11b | 5C6 | FITC | Bio-Rad | MCA711F | Rat IgG2b | 1:100 | <15 | - |
| CD11b | 5C6 | PE | Bio-Rad | MCA711PE | Rat IgG2b | 1:100 | <15 | - |
| CD11b | M1/70 | AF488 | BioLegend | 101217 | Rat IgG2b, κ | 1:100 | <15 | - |
| CD11c | N418 | AF488 | Thermo | MCD11c20 | Hamster IgG | 1:50 | <15 | - |
| CD11c | N418 | AF594 | BioLegend | 117346 | Hamster IgG | 1:50 | >120 | - |
| CD11c | N418 | AF647 | BioLegend | 117312 | Hamster IgG | 1:100 | <15 | - |
| CD11c | B-Ly6 | AF700 | BD Biosciences | 561352 | Mouse $IgG_1$, κ | 1:25 | <15 | - |
| CD20 | L26 | AF488 | Thermo | 53-0202-82 | Mouse IgG2b, κ | 1:200 | <15 | - |
| CD21 | 7E9 | Pacific Blue | BioLegend | 123413 | Rat IgG2a, κ | 1:200 | <15 | - |
| CD21 | Bu32 | AF532 | BioLegend | Custom | Mouse $IgG_1$, κ | 1:400 | <15 | - |
| CD23 | B3B4 | AF647 | BioLegend | 101611 | Rat IgG2a, κ | 1:50 | <15 | - |
| CD23 | EBVCS-5 | AF532 | BioLegend | NA, Custom | Mouse $IgG_1$, κ | 1:25 | <15 | - |
| CD25 | PC61.5 | AF488 | Thermo | 53-0251-82 | Rat IgG1, λ | 1:50 | <15 | - |
| CD25 | M-A251 | AF647 | BioLegend | 356127 | Mouse $IgG_1$, κ | 1:50 | <15 | - |

| | | | | | | | | |
|---|---|---|---|---|---|---|---|---|
| CD31 | MEC13.3 | AF488 | BioLegend | 102514 | Rat IgG2a, κ | 1:100 | <15 | - |
| CD31 | MEC13.3 | PE | BD Biosciences | 553373 | Rat IgG2a, κ | 1:200 | <15 | - |
| CD31 | MEC13.3 | AF594 | BioLegend | 102520 | Rat IgG2a, κ | 1:100 | >120 | - |
| CD31 | MEC13.3 | AF647 | BioLegend | 102516 | Rat IgG2a, κ | 1:100 | <15 | - |
| CD31 | WM59 | AF700 | BioLegend | 303133 | Mouse $IgG_1$, κ | 1:25 | <15 | - |
| CD34 | QBEND/10 | PE | Thermo | MAI-10205 | Mouse $IgG_1$ | 1:50 | <15 | - |
| CD35 | 8C12 | BV510 | BD Biosciences | 740132 | Rat IgG2a, κ | 1:600 | - | <15 |
| CD35 | E11 | PE | BioLegend | 333406 | Mouse $IgG_1$, κ | 1:800 | <15 | - |
| CD38 | HIT2 | AF700 | BioLegend | 303524 | Mouse $IgG_1$, κ | 1:25 | <15 | - |
| CD39 | A1 | PE | BioLegend | 328208 | Mouse $IgG_1$, κ | 1:50 | <15 | - |
| CD44 | IM7 | AF488 | BioLegend | 103016 | Rat IgG2b, κ | 1:100 | <15 | - |
| CD44 | IM7 | AF532 | BioLegend | Custom | Rat IgG2b, κ | 1:100 | <15 | - |
| CD44 | IM7 | AF647 | BioLegend | 103018 | Rat IgG2b, κ | 1:100 | <15 | - |
| CD44 | IM7 | AF700 | BioLegend | 103026 | Rat IgG2b, κ | 1:50 | <15 | - |
| CD45 | 30-F11 | BV421 | BioLegend | 103134 | Rat IgG2b, κ | 1:100 | - | <15 |
| CD45 | 30-F11 | BV510 | BioLegend | 103138 | Rat IgG2b, κ | 1:100 | - | <15 |
| CD45 | 30-F11 | AF488 | BioLegend | 103122 | Rat IgG2b, κ | 1:200 | <15 | - |
| CD45 | 30-F11 | AF532 | Thermo | 58-0451-82 | Rat IgG2b, κ | 1:200 | <15 | - |
| CD45 | 30-F11 | AF647 | BioLegend | 103124 | Rat IgG2b, κ | 1:400 | <15 | - |
| CD45 | 30-F11 | AF700 | BioLegend | 103128 | Rat IgG2b, κ | 1:50 | <15 | - |
| CD45 | HI30 | AF532 | Thermo | 58-0459-41 | Mouse $IgG_1$, κ | 1:50 | <15 | - |
| CD45 | F10-89-4 | PE/iFluor594 | Caprico Biotechnologies | 1016185 | Mouse IgG2a, κ | 1:50 | <15 | - |
| CD49a | TS2/7 | PE | BioLegend | 328304 | Mouse $IgG_1$, κ | 1:50 | <15 | - |
| CD54 | HA58 | AF647 | BioLegend | 353114 | Mouse $IgG_1$, κ | 1:50 | <15 | - |
| CD64 | X54-5/7.1 | AF647 | BioLegend | 139322 | Mouse $IgG_1$, κ | 1:50 | <15 | - |
| CD66b | G10F5 | AF647 | BioLegend | 305110 | Mouse IgM, κ | 1:25 | <15 | - |
| CD68 | FA-11 | BV421 | BioLegend | 137017 | Rat IgG2a | 1:200 | - | <15 |
| CD68 | FA-11 | AF488 | BioLegend | 137011 | Rat IgG2a | 1:50 | <15 | - |
| CD68 | KP1 | AF647 | Santa Cruz | sc-200060 | Mouse $IgG_1$, κ | 1:100 | <15 | - |
| CD69 | - | None | R&D | AF2386 | Goat IgG | 1:50 | - | - |
| CD69 | H1.2F3 | PE | BioLegend | 104508 | Hamster IgG | 1:50 | <15 | - |
| CD69 | FN50 | AF647 | BioLegend | 310918 | Mouse $IgG_1$, κ | 1:100 | <15 | - |
| CD94 | DX22 | PE | BioLegend | 305506 | Mouse $IgG_1$, κ | 1:50 | <15 | - |
| CD103 | 2E7 | AF488 | BioLegend | 121407 | Hamster IgG | 1:100 | <15 | - |
| CD106 | 429 | AF488 | BioLegend | 105710 | Rat IgG2a, κ | 1:50 | <15 | - |
| CD106 | 429 | eF660 | Thermo | 50-1061-80 | Rat IgG2a, κ | 1:50 | <15 | - |
| CD106 | STA | PE | BioLegend | 305806 | Mouse $IgG_1$, κ | 1:50 | <15 | - |
| CD117 | 2B8 | BV421 | BioLegend | 105827 | Rat IgG2b, κ | 1:50 | - | <15 |
| CD117 | 104D2 | AF488 | BioLegend | 313234 | Mouse $IgG_1$, κ | 1:50 | <15 | - |
| CD138 | 281-2 | BV421 | BD Biosciences | 562610 | Rat IgG2a, κ | 1:50 | - | <15 |
| CD138 | 281-2 | AF647 | BioLegend | 142526 | Rat IgG2a, κ | 1:50 | <15 | - |
| CD138 | MI15 | PE | BioLegend | 356504 | Mouse $IgG_1$, κ | 1:200 | <15 | - |
| CD138 | MI15 | AF647 | BioLegend | 356523 | Mouse $IgG_1$, κ | 1:200 | <15 | - |
| CD163 | GH1/61 | AF647 | BioLegend | 333620 | Mouse $IgG_1$, κ | 1:100 | <15 | - |
| CD166 | EPR2759 | AF488 | AbCAM | Ab197543 | Rabbit mAb | 1:100 | <15 | - |
| CD169 | 3D6.112 | FITC | Bio-Rad | MCA884F | Rat IgG2a, κ | 1:50 | <15 | - |
| CD169 | 3D6.112 | PE | Biolegend | 142404 | Rat IgG2a, κ | 1:300 | <15 | - |
| CD169 | 3D6.112 | AF594 | BioLegend | 142416 | Rat IgG2a, κ | 1:100 | >120 | - |
| CD200 | Ox-90 | BV421 | BD Biosciences | 565547 | Rat IgG2a, κ | 1:100 | - | <15 |
| CD206 | C068C2 | BV421 | BioLegend | 141717 | Rat IgG2a, κ | 1:100 | - | <15 |
| CD207 | eBioL31 | PE | Thermo | 12-2075-80 | Rat IgG2a | 1:50 | <15 | - |
| Clec9a | 8F9 | AF488 | BioLegend | Custom | Mouse IgG2a, κ | 1:25 | <15 | - |
| Collagen IV | - | None | AbCAM | 19808 | Rabbit IgG | 1:200 | - | - |
| Collagen IV | - | None | AbCAM | Ab6586 | Rabbit IgG | 1:200 | - | - |
| CXCL12 | 79018 | AF532 | R&D | MAB350-500 (Unconjugated) | Mouse $IgG_1$ | 1:25 | <15 | - |
| CXCL13 | - | AF532 | R&D | AF801 (Unconjugated) | Goat IgG | 1:25 | <15 | - |
| CXCR6 | 221002 | AF647 | Novus | FAB2145R | Rat IgG2b | 1:50 | <15 | - |
| Cytokeratin | C-11 | AF647 | BioLegend | 628604 | Mouse $IgG_1$, κ | 1:200 | <15 | - |
| Cytokeratin | AE1/AE3 | eF660 | Thermo | 50-9003-82 | Mouse $IgG_1$ | 1:100 | <15 | - |
| DCAMKL1 | - | None | AbCAM | Ab37994 | Rabbit IgG | 1:50 | - | - |

| | | | | | | | | |
|---|---|---|---|---|---|---|---|---|
| DC-SIGN | 9E9A8 | AF647 | BioLegend | 330112 | Mouse IgG2a, κ | 1:50 | <15 | - |
| DEC205 | NLDC-145 | AF647 | BioLegend | 138204 | Rat IgG2a, κ | 1:50 | <15 | - |
| Desmin | - | None | AbCAM | Ab15200 | Rabbit IgG | 1:200 | - | - |
| Desmin | Y66 | AF488 | AbCAM | Ab185033 | Rabbit IgG | 1:200 | <15 | - |
| E-cadherin | DECMA-1 | AF647 | BioLegend | 147308 | Rat IgG1, κ | 1:100 | <15 | - |
| EpCAM | G8.8 | BV510 | BD Biosciences | 563216 | Rat IgG2a, κ | 1:100 | - | <15 |
| EpCAM | G8.8 | AF594 | BioLegend | 118222 | Rat IgG2a, κ | 1:400 | >120 | - |
| EpCAM | G8.8 | AF647 | BioLegend | 118212 | Rat IgG2a, κ | 1:200 | <15 | - |
| EpCAM | 9C4 | AF594 | BioLegend | 324228 | Mouse IgG2b, κ | 1:500 | >120 | - |
| EpCAM | 9C4 | AF647 | BioLegend | 324212 | Mouse IgG2b, κ | 1:100 | <15 | - |
| F4/80 | BM8 | BV421 | BioLegend | 123132 | Rat IgG2a, κ | 1:50 | - | <15 |
| F4/80 | BM8 | PE | Thermo | 12-4801-82 | Rat IgG2a, κ | 1:100 | <15 | - |
| F4/80 | BM8 | AF647 | BioLegend | 123122 | Rat IgG2a, κ | 1:50 | <15 | - |
| F4/80 | BM8 | AF700 | BioLegend | 123130 | Rat IgG2a, κ | 1:50 | <15 | - |
| Fibronectin | 2F4 | AF532 | Novus | NBP2-22113AF532 | Mouse $IgG_1$ | 1:25 | <15 | - |
| Foxp3 | FJK-16s | AF488 | Thermo | 53-5773-82 | Rat IgG2a, κ | 1:50 | <15 | - |
| Foxp3 | FJK-16s | AF532 | Thermo | 58-5773-82 | Rat IgG2a, κ | 1:50 | <15 | - |
| Foxp3 | FJK-16s | PE | Thermo | 12-5773-82 | Rat IgG2a, κ | 1:50 | <15 | - |
| Foxp3 | FJK-16s | eF570 | Thermo | 41-5773-82 | Rat IgG2a, κ | 1:50 | <15 | - |
| Foxp3 | FJK-16s | eF660 | Thermo | 50-5773-82 | Rat IgG2a, κ | 1:50 | <15 | - |
| FOXP3 | 236A/E7 | eF570 | Thermo | 41-4777-82 | Mouse $IgG_1$, κ | 1:50 | <15 | - |
| GL-7 | GL7 | PE | BD Biosciences | 561530 | Rat IgM, κ | 1:100 | <15 | - |
| Glutamine synthetase | - | None | AbCAM | Ab49873 | Rabbit IgG | 1:200 | - | - |
| gp38 | 8.1.1 | AF488 | BioLegend | 127405 | Hamster IgG | 1:50 | <15 | - |
| HLA-DR | L243 | AF488 | BioLegend | 307619 | Mouse IgG2a, κ | 1:200 | <15 | - |
| Hoechst | - | - | Biotium | 40046 | - | 1:5,000 | - | - |
| ICOS | CS98.4A | AF488 | BioLegend | 313514 | Hamster IgG | 1:25 | <15 | - |
| IgA | - | AF555 | Southern Biotech | 1040-32 | Goat IgG | 1:500 | <15 | - |
| IgA1 | B3506B4 | AF647 | SouthernBiotech | 9130-31 | Mouse $IgG_1$, κ | 1:500 | <15 | - |
| IgA2 | A9604D2 | AF488 | SouthernBiotech | 9140-30 | Mouse $IgG_1$, κ | 1:500 | <15 | - |
| IgD | 11-26c.2a | AF488 | BioLegend | 405718 | Rat IgG2a, κ | 1:400 | <15 | - |
| IgD | 11-26c.2a | AF594 | BioLegend | 405740 | Rat IgG2a, κ | 1:400 | >120 | - |
| IgD | 11-26c.2a | AF700 | BioLegend | 405729 | Rat IgG2a, κ | 1:50 | <15 | - |
| IgD | IA6-2 | AF488 | BioLegend | 348216 | Mouse IgG2a, κ | 1:25 | <15 | - |
| IgM | EPR5539-65-4 | AF647 | AbCAM | Ab200629 | Rabbit mAb | 1:100 | <15 | - |
| IRF4 | 3E4 | FITC | Thermo | 11-9858-82 | Rat IgG1, κ | 1:50 | <15 | - |
| IRF4 | 3E4 | PE | Thermo | 12-9858-82 | Rat IgG1, κ | 1:50 | <15 | - |
| JOJO-1 | - | - | Thermo | J11372 | - | 1:10,000 | - | - |
| Keratin 14 | Poly9060 | - | BioLegend | 906004 | Chicken IgY | 1:50 | - | - |
| Keratin 18 | 1G11C4 | CoraLite488 | ProteinTech | CL488-66187 | Mouse $IgG_1$ | 1:50 | <15 | - |
| Ki-67 | B56 | AF488 | BD Biosciences | 558616 | Mouse $IgG_1$, κ | 1:50 | <15 | - |
| Ki-67 | B56 | AF700 | BD Biosciences | 561277 | Mouse $IgG_1$, κ | 1:50 | <15 | - |
| KLRG1 | 2F1 | AF488 | BD Biosciences | 561619 | Hamster $IgG_2$, κ | 1:50 | <15 | - |
| Laminin 1 + 2 | - | None | AbCAM | Ab7463 | Rabbit IgG | 1:100 | - | - |
| Lumican | - | AF532 | R&D | AF2846 (Unconjugated) | Goat IgG | 1:50 | <15 | - |
| Ly-6G | 1A8 | AF488 | BioLegend | 127626 | Rat IgG2a, κ | 1:50 | <15 | - |
| Lysozyme | - | - | AbCAM | Ab2408 | Rabbit IgG | 1:50 | - | - |
| Lyve-1 | ALY7 | eF570 | Thermo | 41-0443-82 | Rat IgG1, κ | 1:100 | <15 | - |
| Lyve-1 | - | AF532 | R&D | AF2089 (Unconjugated) | Goat IgG | 1:100 | <15 | - |
| MARCO | - | - | Thermo | PA5-64134 | Rabbit IgG | 1:25 | - | - |
| MHC-II | M5/114.15.2 | BV421 | BioLegend | 107631 | Rat IgG2b, κ | 1:400 | - | <15 |
| MHC-II | M5/114.15.2 | AF647 | BioLegend | 107618 | Rat IgG2b, κ | 1:600 | <15 | - |
| MHC-II | M5/114.15.2 | AF700 | BioLegend | 107622 | Rat IgG2b, κ | 1:100 | <15 | - |
| NF-H/NF-M | SM1-35 | AF488 | BioLegend | 835614 | Mouse $IgG_1$, κ | 1:50 | <15 | - |
| NK1.1 | PK136 | BV421 | BioLegend | 108731 | Mouse IgG2a, κ | 1:50 | - | <15 |
| p53 | PAb 240 | PE | Novus | NB200-103PE | Mouse $IgG_1$, κ | 1:50 | <15 | - |
| Pax5 | 1H9 | AF647 | BioLegend | 649704 | Rat IgG2a, κ | 1:100 | <15 | - |
| PD-1 | 29F.1A.12 | BV421 | BioLegend | 135217 | Rat IgG2a, κ | 1:100 | - | <15 |
| PD-1 | 29F.1A.12 | PE | BioLegend | 135206 | Rat IgG2a, κ | 1:100 | <15 | - |
| PD-1 | EH12.2H7 | PE | BioLegend | 329906 | Mouse $IgG_1$, κ | 1:200 | <15 | - |
| RoRyt | AFKJS-9 | - | Thermo | 14-6988-82 | Rat IgG2a | 1:200 | - | - |

| | | | | | | | | |
|---|---|---|---|---|---|---|---|---|
| SiglecF | E50-2440 | PE | BD Biosciences | 552126 | Rat IgG2a, κ | 1:100 | <15 | - |
| SiglecF | 1RNM44N | AF700 | Thermo | 56-1702-80 | Rat IgG2a, κ | 1:50 | <15 | - |
| SIRPα | P84 | AF488 | BioLegend | 144024 | Rat IgG1, κ | 1:50 | <15 | - |
| SIRPα | P84 | AF647 | BioLegend | 144027 | Rat IgG1, κ | 1:200 | <15 | - |
| αSMA | 1A4 | AF488 | Thermo | 53-9760-80 | Mouse IgG2a, κ | 1:500 | <15 | - |
| αSMA | 1A4 | eF660 | Thermo | 53-9760-82 | Mouse IgG2a, κ | 1:500 | <15 | - |
| SPARC | - | AF532 | R&D | AF941 (Unconjugated) | Goat IgG | 1:50 | <15 | - |
| TCRγδ | GL3 | AF488 | BioLegend | 118128 | Hamster IgG | 1:50 | <15 | - |
| TCRγδ | GL3 | PE | BioLegend | 118108 | Hamster IgG | 1:100 | <15 | - |
| TCRγδ | B1 | PE | BioLegend | 331210 | Mouse $IgG_1$, κ | 1:100 | <15 | - |
| Tim-3 | 344823 | AF532 | R&D | MAB2365 (Unconjugated) | Rat IgG2a | 1:25 | <15 | - |
| Tim-4 | 21H12 | BV421 | BD Biosciences | 742773 | Rat IgG1, κ | 1:100 | - | <15 |
| Tryptase | AA1 | - | AbCAM | Ab2378 | Mouse $IgG_1$ | 1:50 | <15 | - |
| Vα7.2 | 3C10 | AF647 | BioLegend | 351726 | Mouse $IgG_1$, κ | 1:50 | <15 | - |
| Vimentin | O91D3 | AF532 | BioLegend | Custom | Mouse IgG2a | 1:200 | <15 | - |
| anti-chicken IgY | - | FITC | Thermo | SA1-72000 | Donkey IgG | 1:200 | <15 | - |
| anti-goat IgG | - | AF488 | Thermo | A-11055 | Donkey IgG | 1:400 | <15 | - |
| anti-hamster IgG | - | AF647 | Thermo | A-21451 | Goat IgG | 1:400 | <15 | - |
| anti-rabbit IgG | - | AF488 | Thermo | A-11034 | Goat IgG | 1:400 | <15 | - |
| anti-rabbit IgG | - | AF532 | Thermo | A-11009 | Goat IgG | 1:400 | <15 | - |
| anti-rabbit IgG | - | AF555 | Thermo | A-21428 | Goat IgG | 1:400 | <15 | - |
| anti-rabbit IgG | - | AF594 | Thermo | A-11037 | Goat IgG | 1:400 | >120 | - |
| anti-rabbit IgG | - | AF647 | Thermo | A-21245 | Goat IgG | 1:400 | <15 | - |
| anti-rabbit IgG | - | AF700 | Thermo | A-21038 | Goat IgG | 1:400 | <15 | - |
| anti-rabbit IgG | - | AF750 | Thermo | A-21039 | Goat IgG | 1:400 | <15 | - |
| anti-rabbit IgG | - | AF647 | Thermo | A-31573 | Donkey IgG | 1:400 | <15 | - |
| anti-rabbit IgG | - | AF488 | Thermo | Z25302 | Goat $Fab_2$ | - | <15 | - |
| anti-rabbit IgG | - | AF532 | Thermo | Z25303 | Goat $Fab_2$ | - | <15 | - |
| anti-rabbit IgG | - | AF555 | Thermo | Z25305 | Goat $Fab_2$ | - | <15 | - |
| anti-rabbit IgG | - | AF594 | Thermo | Z25307 | Goat $Fab_2$ | - | >120 | - |
| anti-rabbit IgG | - | AF647 | Thermo | Z25308 | Goat $Fab_2$ | - | <15 | - |
| anti-rat IgG | - | AF647 | Thermo | A-21247 | Goat IgG | 1:400 | <15 | - |

### Table S2. IBEX panels for individual organs (See Figs. 3, 4A, and Movies S1-S7).

#### Spleen: 16 parameters, 3 cycles

| Cycle | Marker | Clone | Conjugate | Vendor | Catalog Number | Dilution |
|---|---|---|---|---|---|---|
| 1 | CD8 | 53-6.7 | BV421 | BioLegend | 100738 | 1:200 |
| | Autofluorescence | - | BV510* | - | - | - |
| | Autofluorescence | - | AF488** | - | - | - |
| | B220 | RA3-6B2 | PE | BD Biosciences | 553090 | 1:400 |
| | CD4 | GK1.5 | AF594 | BioLegend | 100446 | 1:200 |
| | Foxp3 | FJK-16s | eF660 | Thermo | 50-5773-82 | 1:50 |
| | IgD | 11-26c.2a | AF700 | BioLegend | 405729 | 1:50 |
| 2 | F4/80 | BM8 | BV421 | BioLegend | 123132 | 1:50 |
| | Autofluorescence | - | BV510* | - | - | - |
| | Collagen IV | - | None | AbCAM | 19808 | 1:200 |
| | Goat anti-rabbit IgG | - | AF488 | Thermo | A-11034 | 1:400 |
| | CD169 | 3D6.112 | PE | Biolegend | 142404 | 1:300 |
| | CD4 | GK1.5 | AF594 | BioLegend | 100446 | 1:200 |
| | CD11c | N418 | AF647 | BioLegend | 117312 | 1:100 |
| | MHC-II | M5/114.15.2 | AF700 | BioLegend | 107622 | 1:100 |
| 3 | CD68 | FA-11 | BV421 | BioLegend | 137017 | 1:200 |
| | Autofluorescence | - | BV510* | - | - | - |
| | CD45 | 30-F11 | AF488 | BioLegend | 103122 | 1:50 |
| | CD31 | MEC13.3 | PE | BD Biosciences | 553373 | 1:200 |
| | JOJO-1 | - | AF532 | Thermo | J11372 | 1:20,000 |
| | CD4 | GK1.5 | AF594 | BioLegend | 100446 | 1:200 |
| | CD3 | 17A2 | AF647 | BD Biosciences | 557869 | 1:400 |
| | Ki-67 | B56 | AF700 | BD Biosciences | 561277 | 1:50 |

#### Thymus: 26 parameters, 5 cycles

| Cycle | Marker | Clone | Conjugate | Vendor | Catalog Number | Dilution |
|---|---|---|---|---|---|---|
| 1 | CD138 | 281-2 | BV421 | BD Biosciences | 562610 | 1:50 |
| | CD11b | 5C6 | FITC | Bio-Rad | MCA711F | 1:100 |
| | β-3 Tubulin | AA10 | AF532 | BioLegend | Custom | 1:100 |
| | B220 | RA3-6B2 | PE | BD Biosciences | 553090 | 1:400 |
| | CD3 | 17A2 | AF594 | BioLegend | 100240 | 1:400 |
| | CD106 | 429 | eF660 | Thermo | 50-1061-80 | 1:50 |
| | Collagen IV | - | None | AbCAM | 19808 | 1:200 |
| | Goat anti-rabbit IgG | - | AF700 | Thermo | A-21038 | 1:400 |
| 2 | CD8 | 53-6.7 | BV421 | BioLegend | 100738 | 1:200 |
| | CD25 | PC61.5 | AF488 | Thermo | 53-0251-82 | 1:50 |
| | Foxp3 | FJK-16s | AF532 | Thermo | 58-5773-82 | 1:50 |
| | CD4 | RM4-5 | eF570 | Thermo | 41-0042-82 | 1:100 |
| | CD3 | 17A2 | AF594 | BioLegend | 100240 | 1:400 |
| | DEC205 | NLDC-145 | AF647 | BioLegend | 138204 | 1:50 |
| | Ki-67 | B56 | AF700 | BD Biosciences | 561277 | 1:50 |
| 3 | CD68 | FA-11 | BV421 | BioLegend | 137017 | 1:200 |
| | SIRPα | P84 | AF488 | BioLegend | 144024 | 1:50 |
| | CD31 | MEC13.3 | PE | BD Biosciences | 553373 | 1:200 |
| | CD3 | 17A2 | AF594 | BioLegend | 100240 | 1:400 |
| | CD11c | N418 | AF647 | BioLegend | 117312 | 1:100 |
| | MHC-II | M5/114.15.2 | AF700 | BioLegend | 107622 | 1:100 |
| 4 | CD206 | C068C2 | BV421 | BioLegend | 141717 | 1:100 |
| | αSMA | 1A4 | AF488 | Thermo | 53-9760-80 | 1:500 |
| | CD44 | IM7 | AF532 | BioLegend | Custom | 1:100 |
| | CD169 | 3D6.112 | PE | Biolegend | 142404 | 1:300 |
| | CD3 | 17A2 | AF594 | BioLegend | 100240 | 1:400 |
| | CD45 | 30-F11 | AF700 | BioLegend | 103128 | 1:50 |
| 5 | JOJO-1 | - | AF532 | Thermo | J11372 | 1:20,000 |
| | TCRγδ | GL3 | PE | BioLegend | 118108 | 1:100 |
| | CD3 | 17A2 | AF594 | BioLegend | 100240 | 1:400 |
| | Cytokeratin | C-11 | AF647 | BioLegend | 628604 | 1:200 |

#### Lung: 23 parameters, 4 cycles

| Cycle | Marker | Clone | Conjugate | Vendor | Catalog Number | Dilution |
|---|---|---|---|---|---|---|
| 1 | CD206 | C068C2 | BV421 | BioLegend | 141717 | 1:100 |
| | CD11b | 5C6 | FITC | Bio-Rad | MCA711F | 1:100 |
| | CD4 | RM4-5 | AF532 | Thermo | 58-0042-80 | 1:50 |
| | Lyve-1 | ALY7 | eF570 | Thermo | 41-0443-82 | 1:100 |

| | | | | | | |
|---|---|---|---|---|---|---|
| | CD31 | MEC13.3 | AF594 | BioLegend | 102520 | 1:100 |
| | CD11c | N418 | AF647 | BioLegend | 117312 | 1:100 |
| | SiglecF | 1RNM44N | AF700 | Thermo | 56-1702-80 | 1:50 |
| 2 | CD68 | FA-11 | BV421 | BioLegend | 137017 | 1:200 |
| | Ly-6G | 1A8 | AF488 | BioLegend | 127626 | 1:50 |
| | β-3 Tubulin | AA10 | AF532 | BioLegend | Custom | 1:100 |
| | B220 | RA3-6B2 | PE | BD Biosciences | 553090 | 1:400 |
| | CD31 | MEC13.3 | AF594 | BioLegend | 102520 | 1:100 |
| | EpCAM | G8.8 | AF647 | BioLegend | 118212 | 1:200 |
| | MHC-II | M5/114.15.2 | AF700 | BioLegend | 107622 | 1:100 |
| 3 | CD138 | 281-2 | BV421 | BD Biosciences | 562610 | 1:50 |
| | KLRG1 | 2F1 | AF488 | BD Biosciences | 561619 | 1:50 |
| | IgA | - | AF555 | Southern Biotech | 1040-32 | 1:500 |
| | CD31 | MEC13.3 | AF594 | BioLegend | 102520 | 1:100 |
| | CD44 | IM7 | AF647 | BioLegend | 103018 | 1:100 |
| | Collagen IV<br>Goat anti-rabbit IgG | -<br>- | None<br>AF700 | AbCAM<br>Thermo | 19808<br>A-21038 | 1:200<br>1:400 |
| 4 | αSMA | 1A4 | AF488 | Thermo | 53-9760-80 | 1:500 |
| | JOJO-1 | - | AF532 | Thermo | J11372 | 1:20,000 |
| | CD169 | 3D6.112 | PE | Biolegend | 142404 | 1:300 |
| | CD31 | MEC13.3 | AF594 | BioLegend | 102520 | 1:100 |
| | CD8 | 53-6.7 | AF647 | BioLegend | 100724 | 1:200 |
| | CD45 | 30-F11 | AF700 | BioLegend | 103128 | 1:50 |

### Small Intestine: 20 parameters, 3 cycles

| Cycle | Marker | Clone | Conjugate | Vendor | Catalog Number | Dilution |
|---|---|---|---|---|---|---|
| 1 | CD8 | 53-6.7 | BV421 | BioLegend | 100738 | 1:200 |
| | CD35 | 8C12 | BV510 | BD Biosciences | 740132 | 1:600 |
| | CD4 | RM4-5 | AF532 | Thermo | 58-0042-80 | 1:50 |
| | Foxp3 | FJK-16s | eF570 | Thermo | 41-5773-82 | 1:50 |
| | EpCAM | G8.8 | AF594 | BioLegend | 118222 | 1:400 |
| | CD3 | 17A2 | AF647 | BD Biosciences | 557869 | 1:400 |
| | IgD | 11-26c.2a | AF700 | BioLegend | 405729 | 1:50 |
| 2 | CD117 | 2B8 | BV421 | BioLegend | 105827 | 1:50 |
| | B220 | RA3-6B2 | BV510 | BioLegend | 103248 | 1:300 |
| | KLRG1 | 2F1 | AF488 | BD Biosciences | 561619 | 1:50 |
| | DCAMKL1<br>Goat anti-rabbit IgG | -<br>- | None<br>AF532 | AbCAM<br>Thermo | Ab37994<br>A-11009 | 1:50<br>1:400 |
| | IgA | - | AF555 | Southern Biotech | 1040-32 | 1:500 |
| | EpCAM | G8.8 | AF594 | BioLegend | 118222 | 1:400 |
| | CD31 | MEC13.3 | AF647 | BioLegend | 102516 | 1:100 |
| | Ki-67 | B56 | AF700 | BD Biosciences | 561277 | 1:50 |
| 3 | CD45 | 30-F11 | BV421 | BioLegend | 103134 | 1:100 |
| | CD11b | 5C6 | FITC | Bio-Rad | MCA711F | 1:100 |
| | JOJO-1 | - | AF532 | Thermo | J11372 | 1:20,000 |
| | Lyve-1 | ALY7 | eF570 | Thermo | 41-0443-82 | 1:100 |
| | EpCAM | G8.8 | AF594 | BioLegend | 118222 | 1:400 |
| | CD11c | N418 | AF647 | BioLegend | 117312 | 1:100 |
| | MHC-II | M5/114.15.2 | AF700 | BioLegend | 107622 | 1:100 |

### Liver: 18 parameters, 4 cycles

| Cycle | Marker | Clone | Conjugate | Vendor | Catalog Number | Dilution |
|---|---|---|---|---|---|---|
| 1 | CD4 | GK1.5 | BV421 | BioLegend | 100443 | 1:200 |
| | Autofluorescence | - | BV510* | - | - | - |
| | Autofluorescence | - | AF488** | - | - | - |
| | LysM-tdTomato* | - | 565-595 nm | - | - | - |
| | Laminin 1 + 2<br>Goat anti-rabbit IgG | -<br>- | None<br>AF594 | AbCAM<br>Thermo | Ab7463<br>A-11037 | 1:100<br>1:400 |
| | E-cadherin | DECMA-1 | AF647 | BioLegend | 147308 | 1:100 |
| 2 | CD8 | 53-6.7 | BV421 | BioLegend | 100738 | 1:200 |
| | Autofluorescence | - | BV510* | - | - | - |
| | Autofluorescence | - | AF488** | - | - | - |
| | Desmin<br>Goat anti-rabbit IgG Fab$_2$ | -<br>- | None<br>AF532 | AbCAM<br>Thermo | Ab15200<br>Z25303 | 1:200<br>- |
| | Laminin 1 + 2<br>Goat anti-rabbit IgG | -<br>- | None<br>AF594 | AbCAM<br>Thermo | Ab7463<br>A-11037 | 1:100<br>1:400 |
| | CXCR6 | 221002 | AF647 | Novus | FAB2145R | 1:50 |
| | CD44 | IM7 | AF700 | BioLegend | 103026 | 1:50 |
| 3 | NK1.1 | PK136 | BV421 | BioLegend | 108731 | 1:50 |
| | Autofluorescence | - | BV510* | - | - | - |
| | Autofluorescence | - | AF488** | - | - | - |
| | CD3 | 17A2 | AF532 | Thermo | 58-0032-80 | 1:50 |
| | Laminin 1 + 2 | - | None | AbCAM | Ab7463 | 1:100 |

| | | | | | | |
|---|---|---|---|---|---|---|
| | Goat anti-rabbit IgG | - | AF594 | Thermo | A-11037 | 1:400 |
| | B220 | RA3-6B2 | AF647 | BioLegend | 103226 | 1:400 |
| | CD45 | 30-F11 | AF700 | BioLegend | 103128 | 1:50 |
| 4 | Tim-4 | 21H12 | BV421 | BD Biosciences | 742773 | 1:100 |
| | Autofluorescence | - | BV510* | - | - | - |
| | CD11b | M1/70 | AF488 | BioLegend | 101217 | 1:100 |
| | Glutamine Synthetase<br>Goat anti-rabbit IgG $Fab_2$ | -<br>- | None<br>AF532 | AbCAM<br>Thermo | Ab49873<br>Z25303 | 1:200<br>- |
| | CD1d | 1B1 | PE | BD Biosciences | 553846 | 1:100 |
| | Laminin 1 + 2<br>Goat anti-rabbit IgG | -<br>- | None<br>AF594 | AbCAM<br>Thermo | Ab7463<br>A-11037 | 1:100<br>1:400 |
| | CD11c | N418 | AF647 | BioLegend | 117312 | 1:100 |
| | MHC-II | M5/114.15.2 | AF700 | BioLegend | 107622 | 1:100 |

*LsyM-tdTomato reporter animal

### Naïve and immunized LNs: 41 parameters, 10 cycles

| Cycle | Marker | Clone | Conjugate | Vendor | Catalog Number | Dilution |
|---|---|---|---|---|---|---|
| 1 | PD-1 | 29F.1.A12 | BV421 | BioLegend | 135218 | 1:100 |
| | CD35 | 8C12 | BV510 | BD Biosciences | 740132 | 1:500 |
| | B220 | RA3-6B2 | AF488 | BD Biosciences | 557669 | 1:400 |
| | JOJO-1 | - | - | Thermo | J11372 | 1:20,000 |
| | CD3 | 17A2 | AF594 | BioLegend | 100240 | 1:100 |
| | Bcl6 | K112-91 | AF647 | BD Biosciences | 561525 | 1:50 |
| | Ki-67 | B56 | AF700 | BD Biosciences | 561277 | 1:50 |
| 2 | CD200 | OX-90 | BV421 | BD Biosciences | 565547 | 1:50 |
| | αSMA | 1A4 | AF488 | Thermo | 53-9760-80 | 1:200 |
| | JOJO-1 | - | - | Thermo | J11372 | 1:20,000 |
| | Lyve-1 | ALY7 | eF570 | Thermo | 41-0443-82 | 1:100 |
| | CD3 | 17A2 | AF594 | BioLegend | 100240 | 1:100 |
| | gp38<br>Goat anti-hamster IgG | 8.1.1<br>- | -<br>AF647 | BioLegend<br>Thermo | 127401<br>A-21451 | 1:50<br>1:400 |
| | Collagen IV<br>Goat anti-rabbit IgG | -<br>- | -<br>AF700 | AbCAM<br>Thermo | 19808<br>A-21038 | 1:200<br>1:400 |
| 3 | F4/80 | BM8 | BV421 | BioLegend | 123132 | 1:50 |
| | CD11b | 5C6 | FITC | Thermo | MA5-16529 | 1:100 |
| | JOJO-1 | - | - | Thermo | J11372 | 1:20,000 |
| | CD207 | eBioL31 | PE | Thermo | 12-2075-80 | 1:50 |
| | CD3 | 17A2 | AF594 | BioLegend | 100240 | 1:100 |
| | CD11c | N418 | AF647 | BioLegend | 117312 | 1:100 |
| 4 | CD8 | 53-6.7 | BV421 | BioLegend | 100738 | 1:200 |
| | JOJO-1 | - | - | Thermo | J11372 | 1:20,000 |
| | CD4 | GK1.5 | PE | BD Biosciences | 553730 | 1:200 |
| | CD3 | 17A2 | AF594 | BioLegend | 100240 | 1:100 |
| | Foxp3 | FJK-16s | eF660 | Thermo | 50-5773-82 | 1:50 |
| 5 | CD206 | C068C2 | BV421 | BioLegend | 141717 | 1:100 |
| | SIRPα | P84 | AF488 | BioLegend | 144024 | 1:50 |
| | JOJO-1 | - | - | Thermo | J11372 | 1:20,000 |
| | TCRγδ | GL3 | PE | BioLegend | 118108 | 1:100 |
| | CD3 | 17A2 | AF594 | BioLegend | 100240 | 1:100 |
| | DEC205 | NLDC-145 | AF647 | BioLegend | 138204 | 1:50 |
| 6 | CD68 | FA-11 | BV421 | BioLegend | 137017 | 1:200 |
| | JOJO-1 | - | - | Thermo | J11372 | 1:20,000 |
| | CD169 | 3D6.112 | PE | Biolegend | 142404 | 1:300 |
| | CD3 | 17A2 | AF594 | BioLegend | 100240 | 1:100 |
| | CD64 | X54-5/7.1 | AF647 | BioLegend | 139322 | 1:50 |
| 7 | CD21 | 7E9 | Pacific Blue | BioLegend | 123413 | 1:200 |
| | CD106 | 429 | AF488 | BioLegend | 105710 | 1:50 |
| | JOJO-1 | - | - | Thermo | J11372 | 1:20,000 |
| | CD31 | MEC13.3 | PE | BD Biosciences | 553373 | 1:200 |
| | CD3 | 17A2 | AF594 | BioLegend | 100240 | 1:100 |
| | CD23 | B3B4 | AF647 | BioLegend | 101611 | 1:50 |
| 8 | CD138 | 281-2 | BV421 | BD Biosciences | 562610 | 1:50 |
| | IRF4 | 3E4 | FITC | Thermo | 11-9858-82 | 1:50 |
| | JOJO-1 | - | - | Thermo | J11372 | 1:20,000 |
| | SiglecF | E50-2440 | PE | BD Biosciences | 552126 | 1:100 |
| | CD3 | 17A2 | AF594 | BioLegend | 100240 | 1:100 |
| | Pax5 | 1H9 | AF647 | BioLegend | 649704 | 1:100 |
| 9 | NK1.1 | PK136 | BV421 | BioLegend | 108731 | 1:50 |
| | KLRG1 | 2F1 | AF488 | BD Biosciences | 561619 | 1:56 |
| | JOJO-1 | - | - | Thermo | J11372 | 1:20,000 |
| | GL-7 | GL7 | PE | BD Biosciences | 561530 | 1:100 |
| | CD3 | 17A2 | AF594 | BioLegend | 100240 | 1:100 |
| | CD44 | IM7 | AF647 | BioLegend | 103018 | 1:100 |
| 10 | MHC-II | M5/114.15.2 | BV421 | BioLegend | 107631 | 1:400 |
| | IgD | 11-26c.2a | AF488 | BioLegend | 405718 | 1:400 |
| | JOJO-1 | - | - | Thermo | J11372 | 1:20,000 |
| | CD3 | 17A2 | AF594 | BioLegend | 100240 | 1:100 |
| | CD45 | 30-F11 | AF647 | BioLegend | 103124 | 1:400 |

**Table S3. IBEX panels for human LNs (See Fig. 5, Movie S8).**

**Metastatic pancreatic LN: 17 parameters, 4 cycles**

| Cycle | Marker | Clone | Conjugate | Vendor | Catalog Number | Dilution |
|---|---|---|---|---|---|---|
| 1 | Hoechst | - | - | Biotium | 40046 | 1:5000 |
| | CD8 | SK1 | AF488 | BioLegend | 344716 | 1:50 |
| | CD3 | UCHT1 | AF532 | Thermo | 58-0038-42 | 1:50 |
| | FOXP3 | 236A/E7 | eF570 | Thermo | 41-4777-82 | 1:50 |
| | EpCAM | 9C4 | AF594 | BioLegend | 324228 | 1:500 |
| | CD25 | M-A251 | AF647 | BioLegend | 356127 | 1:50 |
| | CD4 | RPA-T4 | AF700 | BioLegend | 300526 | 1:25 |
| 2 | Hoechst | - | - | Biotium | 40046 | 1:5000 |
| | CD45 | HI30 | AF532 | Thermo | 58-0459-41 | 1:50 |
| | PD-1 | EH12.2H7 | PE | BioLegend | 329906 | 1:200 |
| | EpCAM | 9C4 | AF594 | BioLegend | 324228 | 1:500 |
| | CD69 | FN50 | AF647 | BioLegend | 310918 | 1:100 |
| 3 | Hoechst | - | - | Biotium | 40046 | 1:5000 |
| | HLA-DR | L243 | AF488 | BioLegend | 307619 | 1:200 |
| | SPARC | Goat IgG | AF532 | R&D | AF941 (Unconjugated) | 1:50 |
| | EpCAM | 9C4 | AF594 | BioLegend | 324228 | 1:500 |
| | CD68 | KP1 | AF647 | Santa Cruz | Sc-200060 | 1:100 |
| | Collagen IV<br>Goat anti-rabbit IgG | -<br>- | None<br>AF700 | AbCAM<br>Thermo | Ab6586<br>A-21038 | 1:200<br>1:400 |
| 4 | Hoechst | - | - | Biotium | 40046 | 1:5000 |
| | CD21 | Bu32 | AF532 | BioLegend | Custom | 1:400 |
| | EpCAM | 9C4 | AF594 | BioLegend | 324228 | 1:500 |
| | CD138 | MI15 | AF647 | BioLegend | 356523 | 1:200 |
| | Ki-67 | B56 | AF700 | BD Biosciences | 561277 | 1:50 |

**Mesenteric LN: 66 parameters, 20 cycles**

| Cycle | Marker | Clone | Conjugate | Vendor | Catalog Number | Dilution |
|---|---|---|---|---|---|---|
| 1 | Hoechst | - | - | Biotium | 40046 | 1:5000 |
| | CD20 | L26 | AF488 | Thermo | 53-0202-82 | 1:200 |
| | SPARC | Goat IgG | AF532 | R&D | AF941 (Unconjugated) | 1:50 |
| | CD10 | FR4D11 | PE | Caprico Biotechnologies | 103926 | 1:50 |
| | CD3 | UCHT1 | AF594 | BioLegend | 300446 | 1:100 |
| | BCL2 | 100 | AF647 | BioLegend | 658705 | 1:25 |
| | Collagen IV<br>Goat anti-rabbit IgG | -<br>- | None<br>AF700 | AbCAM<br>Thermo | Ab6586<br>A-21038 | 1:200<br>1:400 |
| 2 | Hoechst | - | - | Biotium | 40046 | 1:5000 |
| | IgD | IA6-2 | AF488 | BioLegend | 348216 | 1:25 |
| | CD21 | Bu32 | AF532 | BioLegend | NA, Custom | 1:600 |
| | CD138 | MI15 | PE | BioLegend | 356504 | 1:200 |
| | CD3 | UCHT1 | AF594 | BioLegend | 300446 | 1:100 |
| | BCL6 | K112-91 | AF647 | BD Biosciences | 561525 | 1:25 |
| | CD31 | WM59 | AF700 | BioLegend | 303133 | 1:25 |
| 3 | Hoechst | - | - | Biotium | 40046 | 1:5000 |
| | HLA-DR | L243 | AF488 | BioLegend | 307620 | 1:100 |
| | CD23 | EBVCS-5 | AF532 | BioLegend | NA, Custom | 1:25 |
| | CD1c | L161 | PE | BioLegend | 331506 | 1:50 |
| | CD3 | UCHT1 | AF594 | BioLegend | 300446 | 1:100 |
| | CD163 | GH1/61 | AF647 | BioLegend | 333620 | 1:100 |
| | CD11c | B-Ly6 | AF700 | BD Biosciences | 561352 | 1:25 |
| 4 | Hoechst | - | - | Biotium | 40046 | 1:5000 |
| | CD8 | SK1 | AF488 | BioLegend | 344716 | 1:25 |
| | CD4 | RPA-T4 | AF532 | Thermo | 58-0049-42 | 1:25 |
| | FOXP3 | 236A/E7 | eF570 | Thermo | 41-4777-82 | 1:25 |
| | CD3 | UCHT1 | AF594 | BioLegend | 300446 | 1:100 |
| | CD25 | M-A251 | AF647 | BioLegend | 356128 | 1:50 |
| | Ki-67 | B56 | AF700 | BD Biosciences | 561277 | 1:50 |
| 5 | Hoechst | - | - | Biotium | 40046 | 1:5000 |
| | ICOS | CS98.4A | AF488 | BioLegend | 313514 | 1:25 |
| | CXCL13 | Goat IgG | AF532 | R&D | AF801 (Unconjugated) | 1:25 |
| | PD-1 | EH12.2H7 | PE | BioLegend | 329906 | 1:100 |
| | CD3 | UCHT1 | AF594 | BioLegend | 300446 | 1:100 |
| | CD69 | FN50 | AF647 | BioLegend | 310918 | 1:25 |
| 6 | Hoechst | - | - | Biotium | 40046 | 1:5000 |
| | CD117 | 104D2 | AF488 | BioLegend | 313234 | 1:50 |
| | Lyve-1 | Goat IgG | AF532 | R&D | AF2089 (Unconjugated) | 1:100 |

| | | | | | | |
|---|---|---|---|---|---|---|
| | CD35 | E11 | PE | BioLegend | 333406 | 1:800 |
| | CD3 | UCHT1 | AF594 | BioLegend | 300446 | 1:100 |
| | CD68 | KP1 | AF647 | Santa Cruz | sc-20060 | 1:100 |
| | CD38 | HIT2 | AF700 | BioLegend | 303524 | 1:25 |
| 7 | Hoechst | - | - | Biotium | 40046 | 1:5000 |
| | Clec9a | 8F9 | AF488 | BioLegend | Custom | 1:25 |
| | Tim-3 | 344823 | AF532 | R&D | MAB2365<br>(Unconjugated) | 1:25 |
| | IRF4 | IRF4.3E4 | PE | BioLegend | 646404 | 1:25 |
| | CD3 | UCHT1 | AF594 | BioLegend | 300446 | 1:100 |
| | DC-SIGN | 9E9A8 | AF647 | BioLegend | 330112 | 1:50 |
| 8 | Hoechst | - | - | Biotium | 40046 | 1:5000 |
| | CXCL12 | 79018 | AF532 | R&D | MAB350-500<br>(Unconjugated) | 1:25 |
| | TCRγδ | B1 | PE | BioLegend | 331210 | 1:100 |
| | CD3 | UCHT1 | AF594 | BioLegend | 300446 | 1:100 |
| | Vα7.2 | 3C10 | AF647 | BioLegend | 351726 | 1:50 |
| 9 | Hoechst | - | - | Biotium | 40046 | 1:5000 |
| | α-SMA | 1A4 | AF488 | Thermo | 53-9760-82 | 1:100 |
| | CD45 | HI30 | AF532 | Thermo | 58-0459-42 | 1:25 |
| | CD106 | STA | PE | BioLegend | 305806 | 1:50 |
| | CD3 | UCHT1 | AF594 | BioLegend | 300446 | 1:100 |
| | CD44 | IM7 | AF647 | BioLegend | 103018 | 1:50 |
| 10 | Hoechst | - | - | Biotium | 40046 | 1:5000 |
| | Lumican | Goat IgG | AF532 | R&D | AF2745<br>(Unconjugated) | 1:50 |
| | CD34 | QBEND/10 | PE | Thermo | MAI-10205 | 1:50 |
| | CD3 | UCHT1 | AF594 | BioLegend | 300446 | 1:100 |
| | CD54 | HA58 | AF647 | BioLegend | 353114 | 1:50 |
| 11 | Hoechst | - | - | Biotium | 40046 | 1:5000 |
| | NF-H/NF-M | SM1-35 | AF488 | BioLegend | 835614 | 1:50 |
| | CD3 | UCHT1 | AF594 | BioLegend | 300446 | 1:100 |
| | CD66b | G10F5 | AF647 | BioLegend | 305110 | 1:25 |
| 12 | Hoechst | - | - | Biotium | 40046 | 1:5000 |
| | CD166 | EPR2759 | AF488 | AbCAM | Ab197543 | 1:125 |
| | Fibronectin | 2F4 | AF532 | Novus | NBP2-22113AF532 | 1:25 |
| | CD39 | A1 | PE | BioLegend | 328208 | 1:50 |
| | CD3 | UCHT1 | AF594 | BioLegend | 300446 | 1:100 |
| | Cytokeratin | AE1/AE3 | eF660 | Thermo | 50-9003-82 | 1:100 |
| 13 | Hoechst | - | - | Biotium | 40046 | 1:5000 |
| | MARCO<br>Zenon Fab | Rabbit IgG<br>- | -<br>AF532 | Thermo<br>Thermo | PA5-64134<br>Z25303 | 1:25<br>NA |
| | CD94 | DX22 | PE | BioLegend | 305506 | 1:50 |
| | CD3 | UCHT1 | AF594 | BioLegend | 300446 | 1:100 |
| | EpCAM | 9C4 | AF647 | BioLegend | 324212 | 1:100 |
| 14 | Hoechst | - | - | Biotium | 40046 | 1:5000 |
| | β-Tubulin 3 | TUJ1 | AF532 | BioLegend | Custom | 1:50 |
| | CD49a | TS2/7 | PE | BioLegend | 328304 | 1:50 |
| | CD3 | UCHT1 | AF594 | BioLegend | 300446 | 1:100 |
| | IgM | EPR5539-65-4 | AF647 | AbCAM | Ab200629 | 1:100 |
| 15 | Hoechst | - | - | Biotium | 40046 | 1:5000 |
| | IgA2 | A9604D2 | AF488 | SouthernBiotech | 9140-30 | 1:500 |
| | p53 | PAb 240 | PE | Novus | NB200-103PE | 1:50 |
| | CD3 | UCHT1 | AF594 | BioLegend | 300446 | 1:100 |
| | IgA1 | B3506B4 | AF647 | SouthernBiotech | 9130-31 | 1:500 |
| 16 | Hoechst | - | - | Biotium | 40046 | 1:5000 |
| | Laminin 1+2<br>Zenon Fab | Rabbit IgG<br>- | -<br>AF488 | AbCAM<br>Thermo | Ab7463<br>Z25302 | 1:100<br>NA |
| | Lysozyme<br>Zenon Fab | Rabbit IgG<br>- | -<br>AF532 | AbCAM<br>Thermo | Ab2408<br>Z25303 | 1:50<br>NA |
| | CD3 | UCHT1 | AF594 | BioLegend | 300446 | 1:100 |
| 17 | Hoechst | - | - | Biotium | 40046 | 1:5000 |
| | Keratin 18 | 1G11C4 | CoraLite488 | ProteinTech | CL488-66187 | 1:50 |
| | Vimentin | O91D3 | AF532 | BioLegend | Custom | 1:200 |
| | CD3 | UCHT1 | AF594 | BioLegend | 300446 | 1:100 |
| 18 | Hoechst | - | - | Biotium | 40046 | 1:5000 |
| | Desmin | Y66 | AF488 | AbCAM | Ab185033 | 1:200 |
| | CD3 | UCHT1 | AF594 | BioLegend | 300446 | 1:100 |
| | Tryptase<br>Zenon Fab | AA1<br>- | -<br>AF700 | AbCAM<br>Thermo | Ab2378<br>Z25011 | 1:50<br>NA |
| 19 | Hoechst | - | - | Biotium | 40046 | 1:5000 |
| | Donkey anti-goat IgG | - | AF488 | Thermo | A11055 | 1:1000 |
| | CD45 | F10-89-4 | PE/iFluor594 | Caprico<br>Biotechnologies | 1016185 | 1:50 |
| | CD3 | UCHT1 | AF594 | BioLegend | 300446 | 1:100 |
| | Donkey anti-rabbit IgG | - | AF647 | Thermo | A31573 | 1:1000 |
| 20 | Hoechst | - | - | Biotium | 40046 | 1:5000 |
| | Keratin 14<br>Donkey anti-chicken IgY | Poly9060<br>- | -<br>FITC | BioLegend<br>Thermo | 906004<br>SA1-72000 | 1:50<br>1:200 |
| | CD3 | UCHT1 | AF594 | BioLegend | 300446 | 1:100 |

| | | | | | | |
|---|---|---|---|---|---|---|
| | Goat anti-rat IgG | - | AF647 | Thermo | A21247 | 1:1000 |

**Table S4. Reagents used for multi-plex Opal IHC experiments (See Figs. 6B, S7A, and Movie S9).**

| Marker | Clone | Conjugate | Vendor | Cat No. | Isotype | Dilution | Time to bleach with $LiBH_4$ |
|---|---|---|---|---|---|---|---|
| CD3e | D4V8L | None | CST | 99940 | Rabbit IgG | 1:100 | - |
| CD4 | D7D2Z | None | CST | 25229 | Rabbit IgG | 1:100 | - |
| CD8 | D4W2Z | None | CST | 98941 | Rabbit IgG | 1:200 | - |
| CD11c | D1V9Y | None | CST | 97585 | Rabbit IgG | 1:100 | - |
| CD19 | D4V4B | None | CST | 90176 | Rabbit IgG | 1:300 | - |
| CD138 | 281-2 | PE | BioLegend | 142504 | Rat IgG2a, κ | 1:100 | - |
| Collagen IV | - | None | AbCam | 19808 | Rabbit IgG | 1:50 | - |
| CXCL9 | - | None | R&D | AF-492-NA | Goat IgG | 1:50 | - |
| E-cadherin | DECMA-1 | AF488 | Thermo | 53-3249-82 | Rat IgG1, κ | 1:100 | - |
| F4/80 | D2S9R | None | CST | 70076 | Rabbit IgG | 1:100 | - |
| F4/80 | BM8 | BV421 | BioLegend | 123132 | Rat IgG2a, κ | 1:50 | - |
| Foxp3 | D6O8R | None | CST | 12653 | Rabbit IgG | 1:200 | - |
| Laminin | - | None | AbCam | Ab7463 | Rabbit IgG | 1:100 | - |
| MHCII | - | None | AbCam | Ab180779 | Rabbit IgG | 1:100 | - |
| Anti-rabbit IgG | | HRP | | | Goat IgG | 1:5 | - |
| - | - | Opal 520 | Akoya Biosciences | FP1487001KT | - | - | >30 minutes |
| - | - | Opal 540 | Akoya Biosciences | FP1494001KT | - | - | >30 minutes |
| - | - | Opal 570 | Akoya Biosciences | FP1488001KT | - | - | 30 minutes |
| - | - | Opal 620 | Akoya Biosciences | FP1495001KT | - | - | >30 minutes |
| - | - | Opal 650 | Akoya Biosciences | FP1496001KT | - | - | 30 minutes |
| - | - | Opal 690 | Akoya Biosciences | FP1497001KT | - | - | 30 minutes |

**Table S5. Reagents used in combined IBEX oligonucleotide-based staining panel. (See Figs. 6D, S7C-D, and Movie S9).**

| Cycle | Antibody/Dye | Clone | Vendor | Cat No. | Imaging oligo sequence |
|---|---|---|---|---|---|
| 1 | Hoechst | - | Biotium | 40046 | - |
| | SIRPα AF488 | P84 | BioLegend | 144024 | - |
| | Foxp3 AF532 | FJK-16s | Thermo | 58-5773-82 | - |
| | CD31 PE | MEC13.3 | BD Biosciences | 553373 | - |
| | CD11c AF647 | N418 | BioLegend | 117312 | - |
| | Ki-67 AF700 | B56 | BD Biosciences | 561277 | - |
| 2 | Hoechst | - | Biotium | 40046 | - |
| | CD169 AF532* | 3D6.112 | BioLegend | 142425 | ATGACTGTCGTCAATTG |
| | IgD Atto 550* | 11.26c.2a | BioLegend | 405745 | GGACAACGGATATGATG |
| | CD11b AF647* | M1/70 | BioLegend | 101265 | ACAAATGAGCCTTCATG |
| | MHCII IR700* | M5/114.15.2 | BioLegend | 107653 | ATCATACTGGTGACCTG |
| 3 | Hoechst | - | Biotium | 40046 | - |
| | CD45 AF532* | 30-F11 | BioLegend | 103159 | TCTGCTCCATAGCCATG |
| | CD68 Atto 550* | FA-11 | BioLegend | 137031 | TCCCGTGAAAGAAAGTG |
| | CD3 AF647* | 17A2 | BioLegend | 100251 | ATCGAGCGGACATACTG |
| Non-IBEX** | B220 AF488* | RA3-6B2 | BioLegend | 103263 | ATTATGAGGTGTAGGTG |
| | CD11c AF594* | N418 | BioLegend | 117355 | GCAAGCGTCCATAACTG |

*Denotes fluorescent label provided by complementary fluorescent oligonucleotides.

**Additional reagents tested (Figure S7C).

### Movies S1-S9 Legends

**Movie S1. High dimensional imaging of the spleen using IBEX.** Confocal images of mouse spleen tissue from a 3 cycle 16 parameter IBEX experiment with CD4 serving as a fiducial. See Fig. 3. Data are representative of 2 similar experiments.

**Movie S2. High dimensional imaging of the thymus using IBEX.** Confocal images of mouse thymus tissue from a 5 cycle 26 parameter IBEX experiment with CD3 serving as a fiducial. See Fig. 3. Data are representative of 2 similar experiments.

**Movie S3. High dimensional imaging of the lung using IBEX.** Confocal images of mouse lung tissue from a 4 cycle 23 parameter IBEX experiment with CD31 serving as a fiducial. See Fig. 3. Data are representative of 2 similar experiments.

**Movie S4. High dimensional imaging of the small intestine using IBEX.** Confocal images of mouse small intestine tissue from a 3 cycle 20 parameter IBEX experiment with EpCAM serving as a fiducial. See Fig. 3. Data are representative of 2 similar experiments.

**Movie S5. High dimensional imaging of the liver using IBEX.** Confocal images of liver tissue from a LysM-tdTomato mouse. A 4 cycle 18 parameter IBEX experiment was performed with Laminin serving as a fiducial. See Fig. 3. Data are representative of 2 similar experiments.

**Movie S6. High dimensional imaging of naïve and immunized LNs using IBEX.** Confocal images of pLNs from naïve and SRBC-immunized mice from 10 cycle 41 parameter IBEX experiments. See Fig. 4A. Data are representative of 2 similar experiments.

**Movie S7. Comparable staining observed by serial and iterative immunofluorescence methods.** Confocal images of inguinal LN (iLN) or pLNs from SRBC-immunized mice demonstrating qualitatively similar staining patterns when antibody panels were applied on individual sections alone (serial) versus on the same section iteratively (IBEX). See Figs. 4 and S5. Data are representative of 2 similar experiments.

**Movie S8. IBEX scales to capture ultra-high content imaging in large human tissues.** Confocal images of human LN tissue section with metastatic lesions from a 4 cycle 17 parameter IBEX experiment or human mesenteric LN from a 20 cycle 66 parameter IBEX experiment. See Fig. 5. Data are representative of similar experiments in normal and diseased human LNs.

**Movie S9. Extensions of IBEX workflow to include Opal fluorophores and oligo-conjugated antibodies.** Representative confocal images from a 10 parameter 4 cycle IBEX experiment incorporating Opal fluorophores performed on heavily fixed mouse pLN tissue sections. Confocal images from a 13 parameter 3 cycle IBEX experiment performed on mouse inguinal LN sections. Markers were visualized using either fluorescently-conjugated antibodies (Cycle 1) or oligo-conjugated antibodies and complementary fluorescent oligos (Cycles 2-3). See Figs. 6 and S7. Data are representative of 2-4 similar experiments.
